# Enhanced mucosal B- and T-cell responses against SARS-CoV-2 after heterologous intramuscular mRNA prime/intranasal protein boost vaccination with a combination adjuvant

**DOI:** 10.1101/2024.03.28.587260

**Authors:** Gabriel Laghlali, Matthew J. Wiest, Dilara Karadag, Prajakta Warang, Jessica J. O’Konek, Lauren A. Chang, Seokchan Park, Mohammad Farazuddin, Jeffrey J. Landers, Katarzyna W. Janczak, Adolfo García-Sastre, James R. Baker, Pamela T. Wong, Michael Schotsaert

## Abstract

Current COVID-19 mRNA vaccines delivered intramuscularly (IM) induce effective systemic immunity, but with suboptimal immunity at mucosal sites, limiting their ability to impart sterilizing immunity. There is strong interest in rerouting immune responses induced in the periphery by parenteral vaccination to the portal entry site of respiratory viruses, such as SARS-CoV-2, by mucosal vaccination. We previously demonstrated the combination adjuvant, NE/IVT, consisting of a nanoemulsion (NE) and an RNA-based RIG-I agonist (IVT) induces potent systemic and mucosal immune responses in protein-based SARS-CoV-2 vaccines administered intranasally (IN). Herein, we demonstrate priming IM with mRNA followed by heterologous IN boosting with NE/IVT adjuvanted recombinant antigen induces strong mucosal and systemic antibody responses and enhances antigen-specific T cell responses in mucosa-draining lymph nodes compared to IM/IM and IN/IN prime/boost regimens. While all regimens induced cross-neutralizing antibodies against divergent variants and sterilizing immunity in the lungs of challenged mice, mucosal vaccination, either as homologous prime/boost or heterologous IN boost after IM mRNA prime was required to impart sterilizing immunity in the upper respiratory tract. Our data demonstrate the benefit of hybrid regimens whereby strong immune responses primed via IM vaccination are rerouted by IN vaccination to mucosal sites to provide optimal protection to SARS-CoV-2.

## INTRODUCTION

Multiple effective COVID-19 vaccines have been developed which have played a pivotal role in overcoming the acute phase of the pandemic. Some of the most efficacious COVID-19 vaccines have been the mRNA vaccines, which were initially administered intramuscularly (IM) as a prime/boost regimen. However, waning antibody titers and the emergence of antigenically drifted versions of the virus (i.e. Omicron variants) with mutations in the viral spike (S) protein that facilitate immune escape, have limited the duration of vaccine-induced protective immunity.^1–4^ Frequent booster immunizations along with introduction of updated vaccines containing mRNAs encoding for the S proteins of Omicron BA.4/5 and XBB1.5 have been deployed in order to address this waning immunity.^5–8^ These booster vaccines are also given IM and have been shown to enhance circulating B/T cell responses and improve protection from severe disease.^2,9,10^ However, even with these current vaccines and booster regimens, breakthrough infections and viral transmission continue to occur in fully vaccinated individuals, demonstrating that these vaccines do not confer sterilizing immunity.

For respiratory viruses such as SARS-CoV-2, the importance of inducing protective mucosal immune responses, including secretory antibodies and tissue-resident T cells, lies in the potential to block initial infection and viral dissemination to the lower respiratory tract (LRT). Moreover, mucosal immune responses in the respiratory tract play a critical role in preventing viral shedding and transmission and can thereby limit pandemic potential. As such, there has been marked interest in developing improved vaccines which induce robust cross-protective mucosal immunity in addition to systemic immunity. While vaccines against respiratory viruses given via the IM route poorly induce mucosal immune responses, mucosal vaccine delivery and natural infection have both been shown to induce robust mucosal immunity.^11^ Furthermore, hybrid immunity against SARS-CoV-2, the result of vaccine-induced immunity boosted by infection, has been suggested to provide superior protection from re-infection even with antigenically drifted viruses.^12,13^ This is potentially mediated in part through re-routing and boosting vaccine-induced B- and T-cell responses to mucosal sites. Using this paradigm, a similar prime/pull strategy can be employed by utilizing mucosal vaccination to mimic events occurring during natural infection, to boost and re-route existing mRNA vaccine-induced immunity to the respiratory tract. While natural infection results in robust activation of local innate mucosal immune responses, achieving such responses through intranasal (IN) immunization with subunit antigens alone is challenging. However, by using rationally designed adjuvants to target immune receptor pathways activated by viral infection in the mucosa, more tailored immune responses can be induced to potentially confer similar or better outcomes as hybrid immunity.

We previously described a potent adjuvant, NE/IVT, for the mucosal delivery of vaccines.^14–16^ NE/IVT is a combination of an oil-in-water nanoemulsion (NE) and a RIG-I agonist based on an *in vitro* transcribed RNA derived from Sendai virus (strain Cantell) defective interfering RNA (IVT).^17^ NE has established Phase I clinical safety profiles as an IN adjuvant (NCT01354379, NCT04148118).^18^ The NE adjuvant induces mucosal and systemic immune responses mediated at least in part, through TLR2 and 4 activation and through NLRP3 activation via induction of immunogenic apoptosis.^19–21^ IVT is a selective RIG-I agonist and potent inducer of type I interferons (IFN-Is).^17^ As a combined agonist, NE/IVT can thus activate all three major innate receptor classes (TLRs, RLRs, NLRs) necessary for induction of antiviral immune responses.^14^ The NE/IVT adjuvant platform has shown a good safety profile in preclinical models, is compatible with whole virus as well as recombinant protein vaccines, and induces potent systemic and mucosal immune responses when used for intranasal (IN) vaccination.

We report herein that IN recombinant SARS-CoV-2 S protein adjuvanted with NE/IVT can boost and induce (pull) mucosal SARS-CoV-2 spike protein-specific immune responses primed by IM mRNA vaccination with the BNT162b2 vaccine. The heterologous regimen of IM mRNA priming followed by IN NE/IVT/S boost resulted in mucosal IgA responses similar to homologous IN NE/IVT/S prime/boost vaccination, but also induced markedly enhanced T_H_1 polarized T cell responses in the upper respiratory tract draining lymph nodes compared to homologous IM mRNA prime/boost and IN NE/IVT/S prime/boost. The strong mucosal B- and T-cell responses that resulted from this heterologous prime/pull vaccination strategy correlated with optimal cross-protection against various divergent VoCs as reflected in protection from morbidity as well as sterilizing virus control in both the upper (URT) and lower respiratory tracts of experimentally SARS-CoV-2 infected mice. In contrast, IM mRNA prime/boost could not impart sterilizing immunity in the URT. These results highlight the potential of heterologous IN boosting in harnessing the strong systemic immunity imparted by mRNA vaccines to drive potent mucosal immune responses in previously vaccinated individuals.

## RESULTS

### NE/IVT-adjuvanted S protein vaccination as homologous IN prime/boost or heterologous IN boost after IM mRNA prime results in strong serum and mucosal IgG as well as mucosal IgA responses

To profile differences in immune responses induced by homologous versus heterologous prime/boost strategies, C57Bl/6 mice were given two immunizations 4-wks apart (**Fig. 1A**). Mice were primed IM either with BNT162b2 mRNA or PBS, or IN with WT S protein adjuvanted with NE (NE/S) or NE/IVT (NE/IVT/S). Two weeks after the prime (wk2), antibody responses were measured against full-length WT S protein and the receptor binding domain (RBD), as RBD is the major target for virus-neutralizing antibodies. Priming with IM mRNA induced robust S protein-specific total IgG titers that were higher than those induced by priming with IN NE/S (by 0.5 log) or IN NE/IVT/S (by 1 log) (**Fig. 1B**). However, RBD-specific total IgG titers were equivalent between the primed groups (**Fig. 1C**). Mice primed with IM mRNA were then boosted either IM with mRNA, or IN with PBS, S protein alone, NE/S, or NE/IVT/S as indicated in **Fig. 1A**. Additionally, mice primed with IN NE/S were boosted with IN NE/S, and those primed with IN NE/IVT/S were boosted with IN NE/IVT/S as homologous IN/IN prime/boost comparison groups. Finally, to examine the effect of a single IN immunization at the boost timepoint, mice primed with PBS were immunized with IN NE/S or NE/IVT/S. Two weeks post-boost (wk6), serum S- and RBD-specific total IgG titers increased for all groups, and were highest in animals receiving IM mRNA prime/boost and for those which received IM mRNA prime followed by IN NE/S or IN NE/IVT/S boost, which resulted in equivalently high IgG titers (GMT 2.8×10^6^, 2.0×10^6^, 2.8×10^6^ against S protein, respectively). S-specific IgG titers for each of these groups increased by at least two logs after the boost, demonstrating the ability of IN NE/S and NE/IVT/S to boost systemic antibody responses induced by IM mRNA priming as effectively as an additional IM mRNA immunization (**Fig. 1D**). In contrast, IM mRNA prime followed by unadjuvanted IN S protein boost only modestly increased S-specific IgG titers relative to levels induced by IM mRNA prime alone. Mice receiving two homologous IN immunizations with NE/IVT/S mounted comparable serum S-specific IgG titers as the IM mRNA prime/boost group, illustrating the ability to induce strong circulating antibody responses by IN immunization with NE/IVT/S. The inclusion of IVT with NE enhanced the magnitude of the induced S-specific IgG response compared to the singly adjuvanted NE/S group for the homologous IN prime/boost groups, confirming the synergistic activity of the combined NE/IVT adjuvant observed in our previous studies.^14–16^ All singly immunized groups (IM mRNA;IN PBS, IM PBS;IN NE/S, IM PBS;IN NE/IVT/S) induced equivalent S-specific IgG titers. While RBD-specific IgG titers were lower by ∼1log for all groups compared to the S-specific IgG titers, the same relative pattern between treatment groups was maintained (**Fig. 1E**).

**Figure 1.**
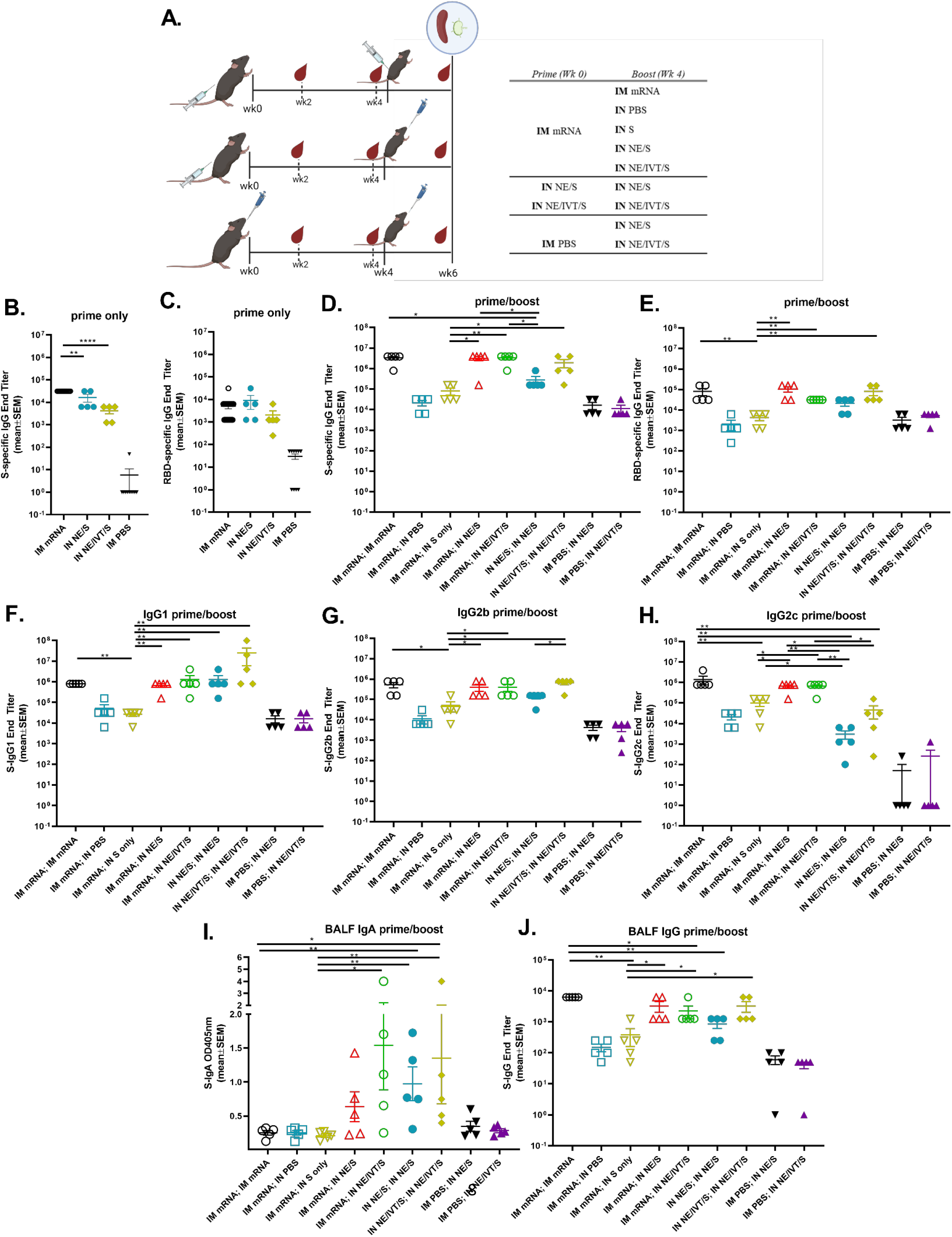
Heterologous IM/IN prime-boost immunization induces robust S protein-specific IgG and enhances mucosal IgA production compared to homologous mRNA IM/IM prime-boost. **(A)** C57Bl/6 mice were given two immunizations 4-wks apart. Mice were primed either IM with 0.25μg of BNT162b2 mRNA or PBS, or IN with 15 μg full-length S protein with either NE or NE/IVT. Mice were then boosted IM with 0.25μg of BNT162b2 mRNA or PBS, or IN with PBS or S protein in PBS, NE or NE/IVT as indicated. Serum antigen-specific total IgG titers against **(B)** WT S protein and **(C)** WT RBD as measured by ELISA 2wks after the prime immunization, and (**D, E**) 2wks after the boost immunization at wk6. (**F-H**) Subclass profiles for S-specific serum antibodies measured at wk6. BALF S-specific (**I**) IgA and (**J**) IgG measured at wk6. (n=5/grp; \**p<0.05*, \*\**p<0.01, ****p<0.0001* by Mann-Whitney U test shown only for select groups-(full statistical analysis is shown in **Table S1**))

IgG subclass skewing was dependent upon the vaccination regimen. IM mRNA prime/boost, heterologous IM mRNA prime followed by IN NE/S or NE/IVT/S boost, as well as IN NE/IVT/S prime/boost all resulted in high and similar antigen-specific IgG1 and IgG2b titers, with the IN NE/IVT/S prime/boost inducing slightly higher levels of IgG1 (**Fig. 1F,G**). The presence of IVT in the IN NE/IVT/S prime/boost group enhanced IgG2b relative to the IN NE/S prime/boost group, consistent with the T_H_1 skewing properties of IVT previously observed.^14–16^ mRNA vaccination, either as homologous prime/boost or as prime followed by IN NE/S or NE/IVT/S boost was required to induce the strongest IgG2c antibody responses, inducing equally high titers (GMT 6×10^5^) in these groups, which was enhanced by ∼1log relative to the group given NE/IVT/S prime/boost via the IN route only (**Fig. 1H**).

While the serum antigen-specific antibody levels and subclass profiles were similar between the homologous IM mRNA prime/boost, and the heterologous IM mRNA;IN NE/S and IM mRNA;IN NE/IVT/S groups, induction of mucosal IgA responses in bronchoalveolar lavage fluid (BALF) was exclusive to groups receiving IN vaccination with NE/S or NE/IVT/S, either as a boost immunization in IM mRNA primed mice, or as a homologous IN prime/boost regimen (**Fig. 1I**). No BALF S-specific IgA was detectable in the IM mRNA prime/boost group. Moreover, no S-specific IgA was observed for mice given IM mRNA;IN S alone, highlighting the role of the NE and NE/IVT adjuvants in driving the mucosal response after IN administration. Interestingly, while a single IN NE/S or NE/IVT/S immunization induced low or no IgA, immunization with IM mRNA;IN NE/IVT/S resulted in equivalent IgA titers as homologous IN NE/IVT/S prime/boost. These results demonstrate the ability of the adjuvanted IN “pull” immunization to harness parenterally primed immune responses for driving robust mucosal immune responses. While dimeric IgA plays the predominant role in first-line defense in the mucosa of the upper respiratory tract, mucosal IgG also contributes to protection through transudation from the blood into the lung^22^. Indeed, in contrast to the pattern observed for IgA, BALF IgG correlated with serum IgG titers, with IM mRNA prime/boost immunization inducing robust BALF antigen-specific IgG titers similar to the IM mRNA;IN NE/S and IM mRNA;IN NE/IVT/S treatment groups, as well as the IN NE/IVT/S prime/boost group (**Fig. 1J**).

### Cross-reactive serum neutralizing antibody titers reflect serum S-specific IgG binding antibody titers

We next quantified cross-reactive neutralizing antibody (nAb) titers induced in immunized C57Bl/6 mice using pseudotyped viruses carrying the S proteins of SARS-CoV-2 variants of concern (VoCs). In general, vaccination regimens that resulted in the highest IgG binding titers (IM mRNA prime/boost, IN NE/S and NE/IVT/S prime/boost or heterologous IM mRNA;IN NE/S or IM mRNA;IN NE/IVT/S) not only resulted in the highest nAb titers against vaccine matched ancestral virus (**Fig. 2A**), but also against the antigenically more distant B.1.617.2 (Delta, **Fig. 2B**), and B.1.351 (Beta, **Fig. 2C**) variants. These treatment groups induced similar levels of cross-neutralizing nAbs against each variant examined. While IM mRNA;IN NE/S immunization induced slightly higher nAbs against the WT and B.1.351 variants relative to homologous IM mRNA or homologous IN NE/S prime/boost, homologous IM mRNA and homologous IN NE/IVT/S prime/boost induced similar titers as heterologous IM mRNA;IN NE/IVT/S immunization. In contrast, IM mRNA primed mice boosted IN with unadjuvanted S alone induced the lowest nAb titers, giving similar or lower titers as the single IM mRNA immunization group against all four variants tested. Neutralization potential was reduced across all vaccination groups by a similar degree (∼2log) towards the more antigenically distant B.1.1.529 (Omicron BA.1) variant, maintaining the same relative pattern of nAb response magnitude observed between immunization groups as against the WT virus (**Fig. 2D**)

**Figure 2:**
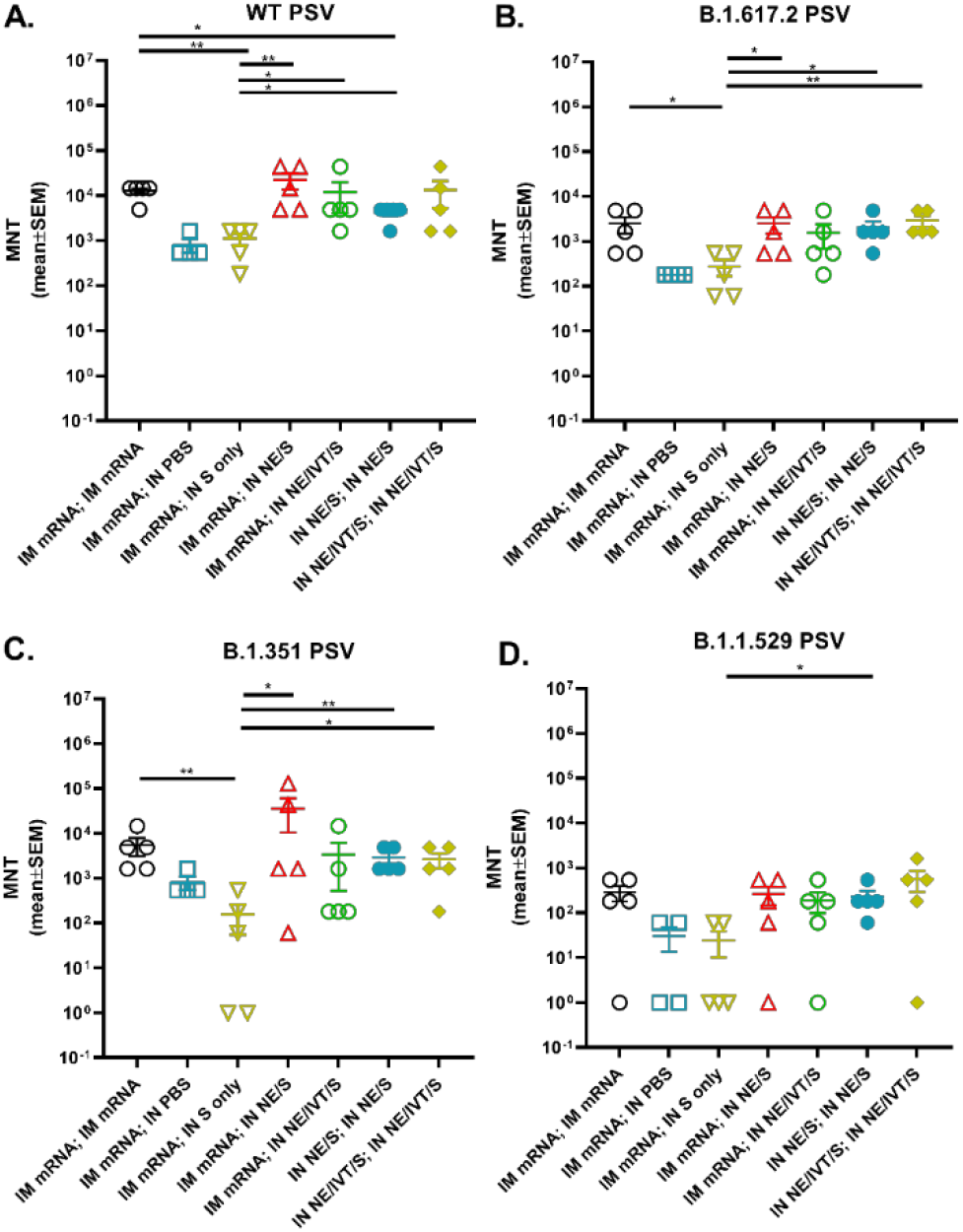
Heterologous IM/IN prime-boost immunization induces robust cross-neutralizing antibody responses against multiple variant viruses. Serum neutralizing antibody titers from immunized C57Bl/6 mice at wk 6 after both prime/boost immunizations with the indicated adjuvant/antigen regimens were measured using pseudoviruses for the **(A)** wild-type, **(B)** B.1.617.2, **(C)** B.1.351, and **(D)** B.1.1.529 (BA.1) variants. (n=5/grp; \**p<0.05*, \*\**p<0.01* by Mann-Whitney U test shown only for select groups-(full statistical analysis is shown in **Table S1**))

### Heterologous IM mRNA prime followed by IN NE/IVT/S boost markedly enhances T_H_1 and T_H_17 polarized antigen recall responses in spleen and cervical lymph nodes

To evaluate T cell antigen recall responses, 2 wks after the boost immunization, splenocytes and cervical lymph node (cLN) isolates were harvested from immunized mice and restimulated *ex vivo* with S protein. Heterologous boosting of IM mRNA primed mice with IN NE/S or IN NE/IVT/S resulted in a marked enhancement of T_H_1-polarized responses compared to homologous IM mRNA, IN NE/S, or IN NE/IVT/S prime/boost groups. High levels of S-specific IFN-γ were induced in splenocytes by IM mRNA vaccination boosted with either IM mRNA or IN NE/S or NE/IVT/S, with the heterologous IN boosted groups inducing equivalent (or higher) levels of IFN-γ than the IM mRNA prime/boost group. The heterologous IM mRNA;IN NE/S and IM mRNA;IN NE/IVT/S groups also demonstrated enhanced levels of IFN-γ compared to the homologous IN NE/S and IN NE/IVT/S prime/boost groups (**Fig. 3A**). As seen previously, inclusion of IVT in the IN NE/IVT/S prime/boost significantly enhanced the IFN-γ response relative to IN NE/S (**Fig. 3A**). Similar patterns were observed for antigen-specific IL-2 responses, with enhanced cytokine levels in animals primed with IM mRNA and then boosted with IN NE/S or NE/IVT/S compared to those given two doses of IM mRNA, IN NE/S, or IN NE/IVT/S (**Fig. 3B**). While significant variation was observed for IP-10, IM mRNA primed animals boosted with IN NE/S or NE/IVT/S induced similar levels of IP-10 in splenocytes as the IM mRNA prime/boost group, with the exception of one mouse in each of the heterologous groups showing much higher IP-10 (5-10 fold relative to the IM mRNA prime/boost) (**Fig. 3C**). Interestingly, singly immunized groups (IM mRNA;IN PBS, IN PBS;IN NE/S, IN PBS;IN NE/IVT/S) induced higher IP-10 than the corresponding homologous prime/boost groups. No significant differences were observed between vaccination groups for TNF-α in stimulated splenocytes, with all immunized groups inducing similar levels of TNF-α, with the singly immunized IN NE/S and NE/IVT/S groups exhibiting slightly reduced levels compared to their respective prime/boost groups (**Fig. 3D**).

**Figure 3:**
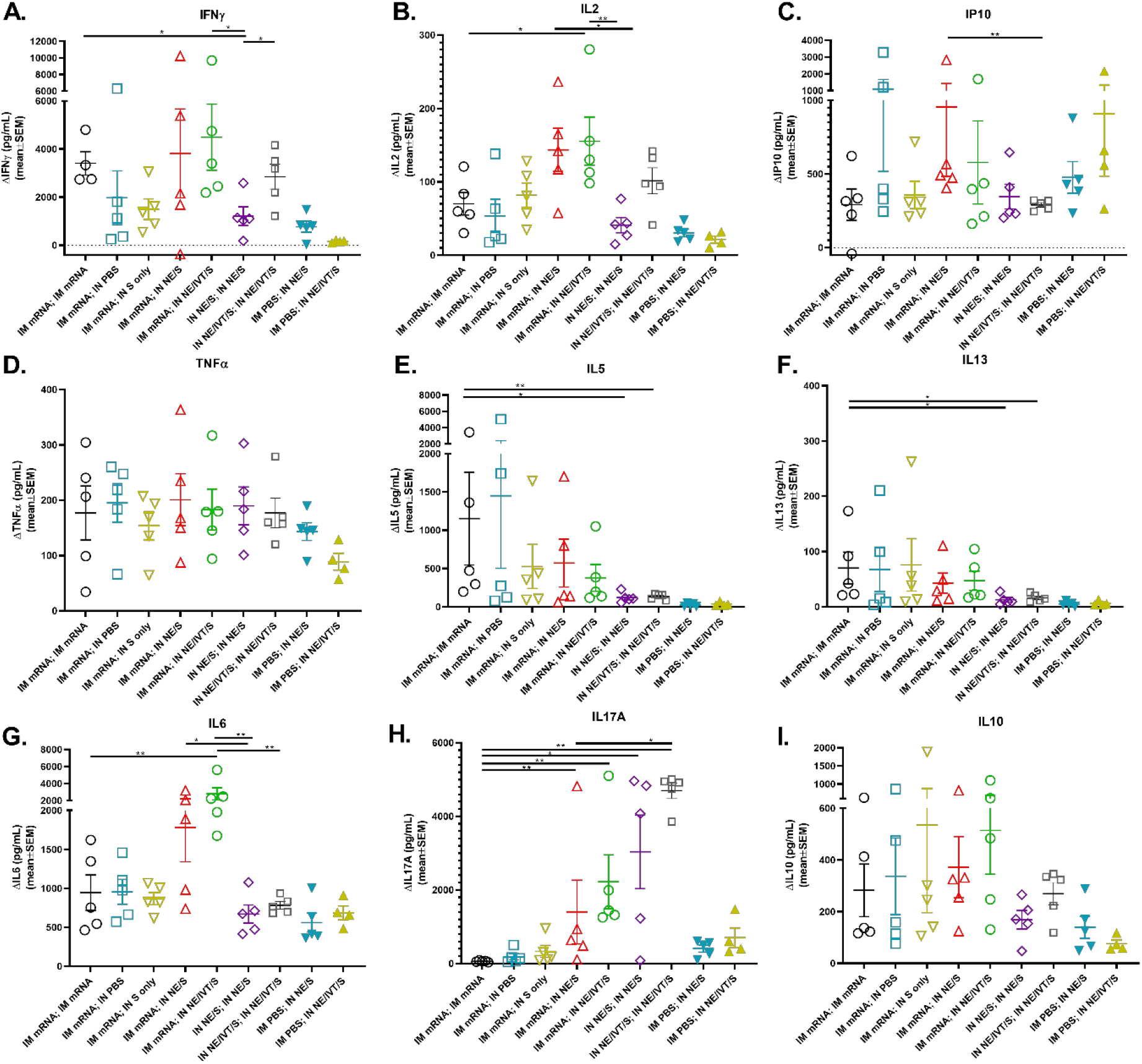
Antigen recall responses assessed in splenocytes isolated from IM/IN immunized mice demonstrate enhanced T_H_1/T_H_17 profiles. Splenocytes were isolated from mice given prime/boost immunizations with the indicated adjuvant/antigen regimens two weeks after the final immunization (wk 6). Splenocytes were stimulated *ex vivo* with 5 μg S protein for 72h, and levels of secreted (A) IFN-γ, (B) IL-2, (C) IP-10, (D) TNF-α, (E) IL-5, (F) IL-13, (G) IL-6, (H) IL-17A, and (I) IL-10 were measured by multiplex immunoassay relative to unstimulated cells. (n=4-5/grp; \**p<0.05*, \*\**p<0.01* by Mann-Whitney U test shown only for select groups-(full statistical analysis is shown in **Table S1**)).

Homologous IM mRNA administration induced significant T_H_2 responses in splenocytes as measured by IL-5 (**Fig. 3E**) and IL-13 (**Fig. 3F**) in comparison to homologous IN NE/S or IN NE/IVT/S, which did not induce detectable levels of these cytokines. Heterologous boosting of IM mRNA primed animals with IN NE/S or NE/IVT/S did not enhance these T_H_2 cytokines and appeared to reduce IL-5 levels compared to the groups given one or two doses of IM mRNA. Finally, while IM mRNA prime/boost induced substantial IL-6 in the spleen (greater than IN NE/S or NE/IVT/S prime/boost), priming with IM mRNA followed by boosting with IN NE/S or NE/IVT/S resulted in significantly increased IL-6 production especially with the NE/IVT/S pull (**Fig. 3G**).

We have established through multiple studies the robust induction of IL-17A production by IN administration of the NE adjuvant, which is further enhanced by the inclusion of IVT.^14–16^ This induced T_H_17 response is exclusive to the mucosal route of administration of these adjuvants. Indeed, we observed high levels of IL-17A induction in splenocytes for the IN NE/S prime/boost group, which was further enhanced ∼2-fold in the IN NE/IVT/S prime/boost group (**Fig. 3H**). In contrast, no IL-17A was observed for the IM mRNA prime/boost group. Interestingly, however, while the single IN immunization groups (IM PBS;IN NE/S, IM PBS;IN NE/IVT/S) induced only low levels of IL-17A (∼10-fold lower than the corresponding prime/boost groups), boosting IM mRNA primed animals with IN NE/S or NE/IVT/S induced high levels of IL-17A, demonstrating the ability of the IN adjuvants to boost and shape immune responses primed by the initial IM mRNA vaccination, promoting a shift in IM mRNA-primed T cell responses towards T_Η_17. Notably, IM mRNA boosted with IN S alone did not induce significant IL-17A production. These results are significant, as T_Η_17 responses have been shown to be a critical component of host defense at mucosal sites.^23–25^ While IL-17A induction has been associated with immune pathology in certain contexts, it has been shown to be non-pathogenic in the context of IL-10 co-production.^26,27^ Indeed, all prime/boost immunization groups with an IM mRNA prime induced similarly high levels of IL-10 in the spleen, which was enhanced relative to the single or two dose IN NE/S or IN NE/IVT/S regimens (**Fig. 3I**). The lack of immune pathology was confirmed in our prior studies and the challenge studies discussed below.

Cervical lymph nodes (cLNs) drain the URT, and therefore are relevant for assessment of local protective immunity near the portal entry sites of pulmonary pathogens and for evaluating mucosal T cell responses induced by IN immunization. IN administration of vaccines as part of a boost regimen enhanced S protein-specific cytokine responses within the cLNs, confirming effective pulling of antigen-specific immune responses to mucosal sites (**Fig. 4**). In general, cytokine profiles in the cLN reflected the recall responses measured in spleen but were heavily skewed by the IN vaccination regimens, resulting in greater magnification of the T_H_1/T_H_17 polarization. Boosting with IN NE/IVT/S after IM mRNA prime induced markedly high antigen-specific IFN-γ levels within the cLN (8514±794 pg/mL (mean±SEM)) compared to homologous prime/boost with IM mRNA (667±581 pg/mL), IN NE/S (303±208 pg/mL), or IN NE/IVT/S (2224±814 pg/mL) (**Fig. 4A**). Furthermore, administration of IN NE, or IN NE/IVT formulations either as a homologous prime/boost regimen or as part of a heterologous boost regimen after IM mRNA priming also significantly enhanced antigen-specific IL-2 and IP-10 responses within the cLN compared to two doses of IM mRNA, with the highest levels observed with heterologous IM mRNA;IN NE/IVT/S vaccination (**Fig. 4B, C**). Notably, IM mRNA prime/boost induced only low levels of these cytokines in the cLN. Finally, enhancement in TNF-α was also observed in the cLN for the IM mRNA;IN NE/IVT/S group as compared to the IM mRNA and IN NE/IVT/S prime/boost groups (**Fig. 4D**). These results are of significance, as the co-production of IFN-γ, IL-2, and TNF-α has been established as a strong mediator of optimal control of viral infection and a major correlate of vaccine protection. Furthermore, these results clearly demonstrate the ability of the IN NE/IVT/S boost to drive enhanced T_H_1 polarized T cell responses in the mucosal lymphoid tissue through ‘pulling’ of responses systemically primed by the IM mRNA.

**Figure 4:**
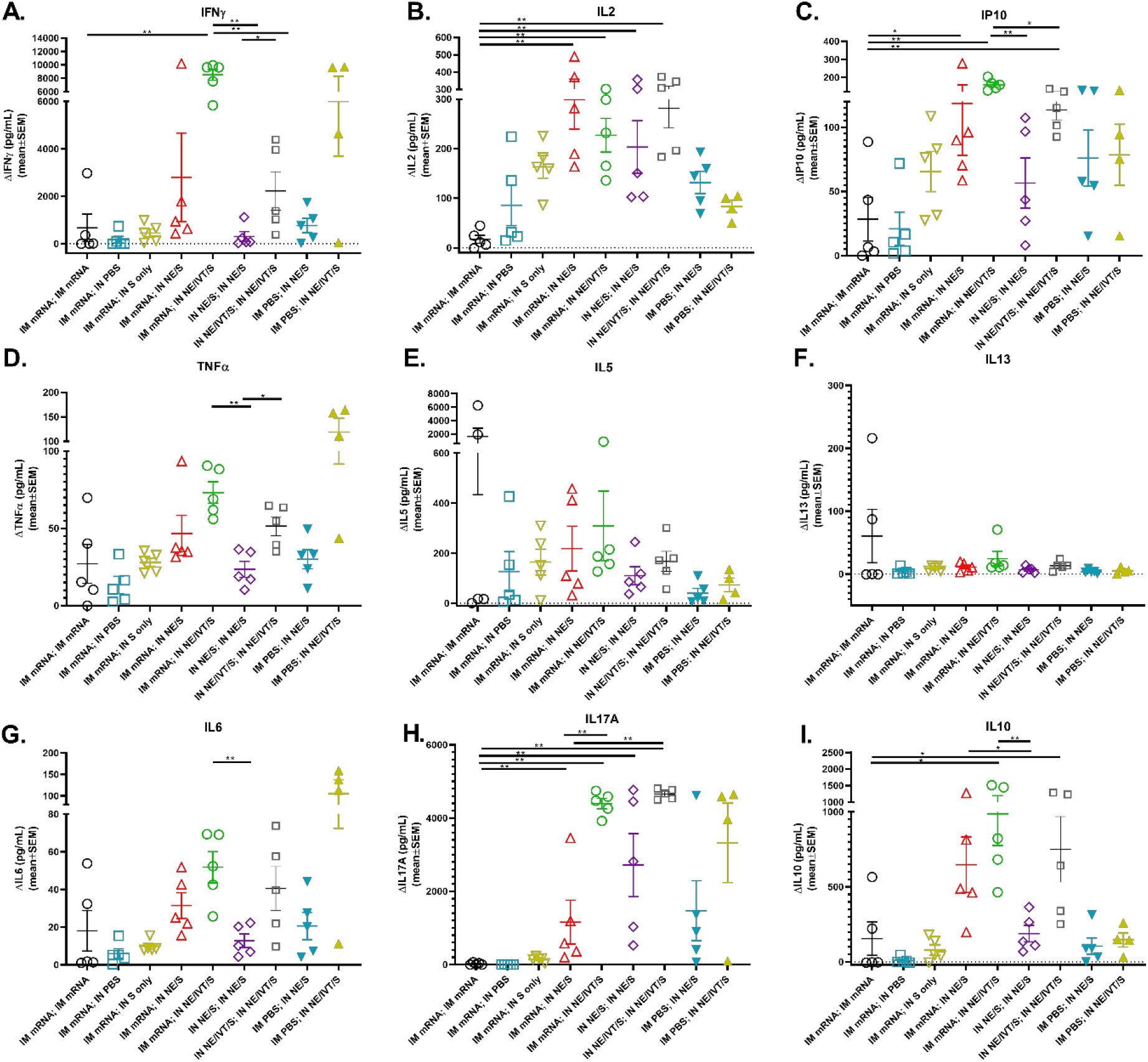
Antigen recall responses assessed in cervical lymph node isolates from IM/IN immunized mice demonstrate even greater enhancement in T_H_1/T_H_17 profiles. cLN cellular isolates from mice given prime/boost immunizations with the indicated adjuvant/antigen regimens were harvested at wk 6 and stimulated *ex vivo* with 5 μg S protein for 72h. Levels of secreted (A) IFN-γ, (B) IL-2, (C) IP-10, (D) TNF-α, (E) IL-5, (F) IL-13, (G) IL-6, (H) IL-17A, and (I) IL-10 were measured by multiplex immunoassay relative to unstimulated cells. (n=4-5/grp; \**p<0.05*, \*\**p<0.01* by Mann-Whitney U test shown only for select groups-(full statistical analysis is shown in **Table S1**)).

Similar to the T_H_2 cytokine pattern in the spleen, two doses of IM mRNA induced the highest levels of IL-5 and IL-13 in the cLN, while two doses of IN NE/S or IN NE/IVT/S induced only low levels of IL-5 and no detectable IL-13 (**Fig. 4E-F**). While heterologous boosting of IM mRNA primed animals with IN NE/S or NE/IVT/S resulted in higher levels of IL-5 than the homologous IN NE/S and IN NE/IVT/S prime/boost groups, the levels remained similar to or lower than those induced in the IM mRNA prime/boost group, and no significant IL-13 was detected in the heterologous prime/boost groups. These results confirm the lack of T_H_2 response enhancement with the adjuvanted heterologous pull immunizations. Minimal levels of spike-specific IL-6 were induced in the cLNs (**Fig. 4G**) as compared to splenocytes (**Fig. 3G**) for all vaccination groups (2 orders of magnitude lower), with the highest levels induced in animals receiving IN NE/IVT/S as part of their vaccine regimen.

The enhancement in IL-17A production seen in the splenocytes was even more pronounced in the cLN for the heterologous IM mRNA;IN NE/IVT/S group (**Fig. 4H**). The combination of IM mRNA prime with IN NE/IVT/S boost resulted in similarly strong IL-17A responses (4398±142 pg/mL) as homologous IN NE/IVT/S prime/boost immunization (4657±136 pg/mL). In contrast, IM mRNA prime/boost did not induce detectable IL-17A. Heterologous boost with IN NE/S was also able to enhance IL-17A in the IM mRNA primed mice, however, inclusion of IVT in the IN boost was critical to driving maximal T_H_17 responses in the cLN. IL-10 production in the cLN followed the same pattern as IL-17A, with the IN adjuvanted groups inducing the highest levels of IL-10 (**Fig. 4I**). Overall, IM mRNA vaccination resulted in priming events that, when boosted IN with NE/IVT/S resulted in a unique antigen-specific cytokine profile in both the spleen and in the local mucosal-draining LNs. These results demonstrate that employing an intranasal “pull” with a NE/IVT adjuvanted vaccine after IM mRNA priming can drive more robust and tailored response towards SARS-CoV-2 through enhancing the T_H_1/T_H_17 polarization.

The induction of high IgG2a, IgG2c, and IgA antibody titers requires efficient class switching in germinal center reactions which require strong CD4^+^ T cell responses. Therefore, we further characterized vaccine induced CD4^+^ T cell responses in the spleens, cLNs, and lungs of immunized animals. To better distinguish differences in cytokine production, C57Bl/6 mice were given the same prime/boost regimens but at higher doses of mRNA (2 μg) and S protein (20 μg) (**Fig. 5**). Mice receiving IM mRNA prime/boost showed robust antigen-specific CD4^+^ T cell responses in the spleen, giving the highest frequencies of IFN-γ^+^, IL-2^+^ and TNF-α^+^ CD4^+^ T cells (**Fig. 5A-C**), as well as polyfunctional (IFN-γ^+^IL-2^+^ TNF-α^+^) CD4^+^ T cells upon stimulation with S protein (**Fig. 5D**). IN boost with S, NE/S or NE/IVT/S after mRNA priming induced similar frequencies of IFN-γ^+^, IL-2^+^ and TNF-α^+^CD4^+^ T cells, as well as polyfunctional CD4^+^ T cells, although at a lower frequency than two doses of IM mRNA. Interestingly, comparison of CD4^+^ IFN-γ^+^ responses by mean fluorescence intensity (MFI) rather than by frequency, showed equivalent, or higher MFIs for the IM mRNA prime with IN S, NE/S, or NE/IVT/S boost groups as the IM mRNA prime/boost group, demonstrating that even though the frequency of IFN-γ^+^ cells was slightly lower for the IM/IN groups, these cells produced higher levels of IFN-γ (**Fig. S3**). In the cLN, however, IM mRNA groups receiving IN NE/S or NE/IVT/S boost developed the strongest antigen-specific CD4^+^T cell responses, with significant enhancement in the frequency of polyfunctional cells compared to groups that received a second IM mRNA dose or IN S (**Fig. 5E-H**). While IN NE/S or NE/IVT/S in a prime/boost regimen induced IL-2 and TNF-α expressing CD4^+^ T cells, IM mRNA priming was necessary to induce optimal IFN-γ expressing CD4^+^ T cells in the cLN, demonstrating the role of the initial IM prime immunization in driving optimal local mucosal responses after a mucosal boost. Furthermore, while IN S boost of IM mRNA primed mice showed enhanced polyfunctional CD4^+^ T cell responses in the spleen comparable to the IN NE/S and NE/IVT/S boost groups, the unadjuvanted boost group showed minimal levels of polyfunctional CD4^+^ T cells in the cLNs, highlighting the critical role of the NE-based adjuvants in driving robust local mucosal cellular responses. Similar to the cLN, in the lungs, groups primed with IM mRNA and then boosted IN with S, NE/S, or NE/IVT/S had similar levels of IFN-γ expressing CD4^+^ T cells which were enhanced compared to the other vaccination regimens (**Fig. 5I-L**). These groups also displayed enhanced polyfunctional CD4^+^ T cells in the lung compared to the other vaccination regimens. Interestingly, boosting with IN S or NE/S after IM mRNA priming induced higher frequencies of polyfunctional CD4^+^ T cells in the lung compared to boosting with IN NE/IVT/S, which was the reverse pattern observed in the cLN, potentially suggesting differences in trafficking and kinetics at this time point between the free antigen vs. NE/IVT adjuvanted S protein. Finally, in accordance with the cytokine secretion data, mucosal boost of IM mRNA primed animals displayed higher frequencies of IL-17A^+^CD4^+^ T cells in the lung than the IM mRNA prime/boost group which had no detectable response (**Fig. S4**). However, the highest frequencies of IL-17A^+^CD4^+^ T cells were induced in the IN NE/S and IN NE/IVT/S homologous prime/boost groups. Taken together, these results demonstrate that boosting with a IN NE antigen regimen represents an effective pulling strategy to mucosal sites of systemic immune responses induced by mRNA vaccination.

**Figure 5:**
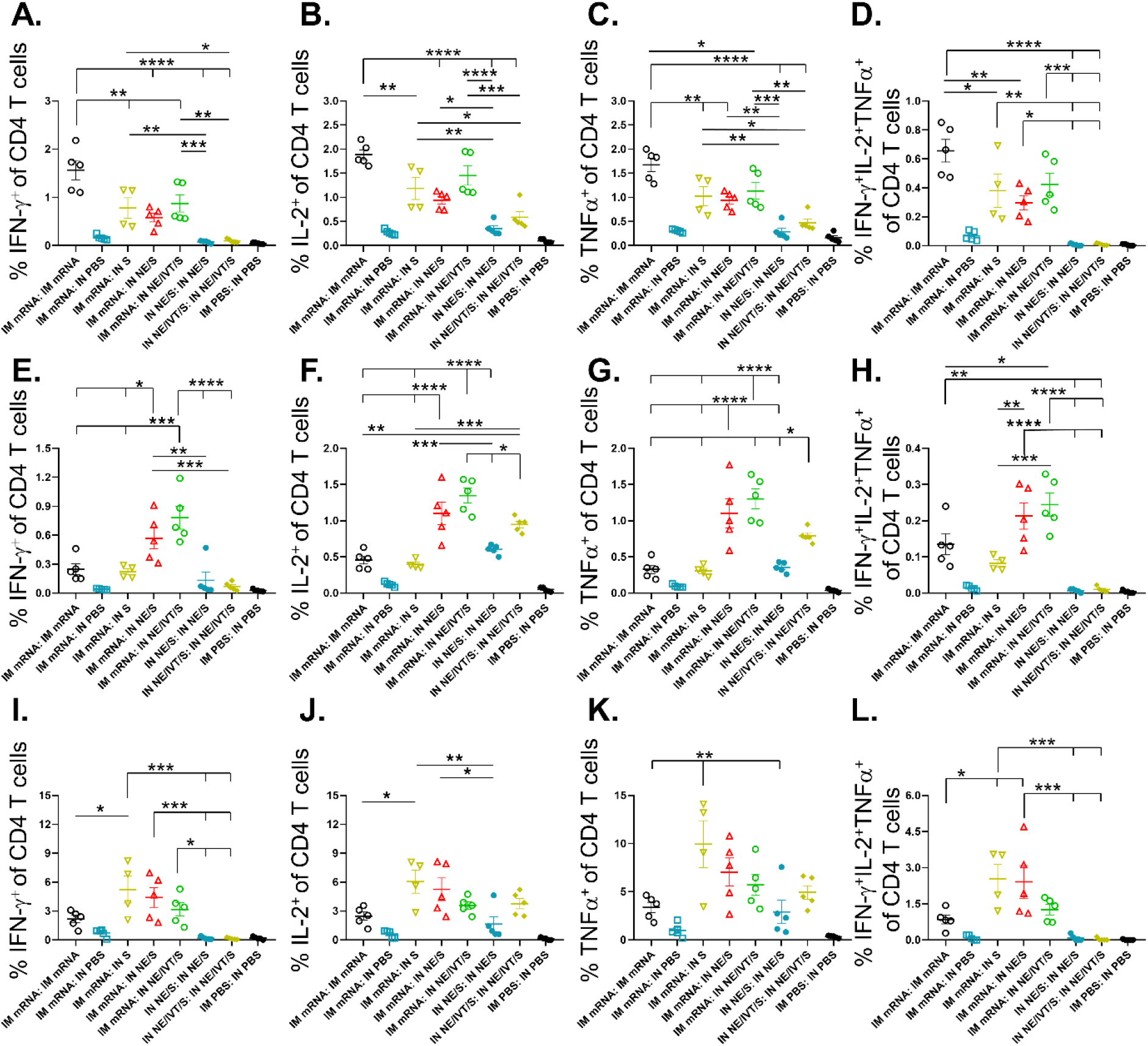
IN administration of antigen with NE adjuvants after IM mRNA priming effectively pulls antigen-specific immunity to mucosal sites. Single cell suspension were isolated from the spleen, cLNs, and lungs of mice given prime/boost immunizations with the indicated adjuvant/antigen regimens. Mice were given 2μg mRNA IM, and 20μg of S protein IN in either PBS, NE, or NE/IVT. Cells were stimulated with 25μg/mL of S protein, and antigen-specific cytokine responses were quantified in CD4^+^ T cells by intracellular cytokine staining and FACS analysis. (n=4-5/grp with data represented as mean ± SEM; \**p<0.05*, \*\**p<0.01,* \*\*\**p<0.001,* \*\*\*\**p<0.0001* by one-way ANOVA with Tukey post-hoc test shown only for select groups-(full statistical analysis is shown in **Table S1**)).

### Heterologous IM/IN prime-boost immunization induces robust virus-neutralizing antibody titers in 129S1 and K18-hACE2 mice and results in superior protection in the upper respiratory tract

To examine the effects of genetic background on induced immune responses, we next repeated vaccination with the heterologous prime/boost regimens in 129S1 and K18-hACE2 mice. The 129S1 strain was selected for vaccination/challenge studies as they are WT mice that have shown to be more susceptible to morbidity after experimental infection with SARS-CoV-2 viruses with the N501Y mutation, such as B.1.351.^28^ In contrast, SARS-CoV-2 Omicron lineages do not efficiently infect WT mice but can replicate in the respiratory tract of transgenic K18-hACE2 mice, which overexpress human ACE2 receptor in the epithelia. Thus, we employed this transgenic model for protective efficacy studies with Omicron BA.5 challenge. Prime immunization with IM mRNA, IN NE/S, IN NE/IVT/S induced equivalent S-specific serum IgG in 129S1 mice, giving similar titers as primed C57Bl/6 and K18-hACE2 mice (**Figs. 1B and S1**). Boosting of IM mRNA primed animals with IM mRNA, IN NE/S or IN NE/IVT/S resulted in high serum total binding IgG against WT S protein in both genetic backgrounds (**Figs. 6A, F**). In both 129S1 and K18-hACE2 mice, comparable induction of high S-specific IgG titers with the IM mRNA prime/boost was observed as for the heterologous IM mRNA prime with IN NE/S or NE/IVT/S boost, with these titers being enhanced compared to the IN NE/S and IN NE/IVT/S prime/boost, and IM mRNA;IN S groups. Overall, an identical pattern for S-specific IgG titers between immunization groups was observed for the 129S1, K18-hACE2, and C57Bl/6 mice. Furthermore, for comparison we assessed whether prime/boost immunization with S protein adjuvanted with the IM adjuvant, Addavax (Advx), would induce similar immune responses. IM Advx/S prime/boost resulted in similar S-specific IgG as the IM mRNA prime/boost group in both 129S1 and K18-hACE2 strains (**Fig 6A, F**). Interestingly, while IM mRNA;IN NE/S and IM mRNA;IN NE/IVT/S groups induced similarly robust neutralization titers as the IM mRNA prime/boost groups in 129S1, K18-hACE2, and C57Bl/6 mice against ancestral virus, a more distinct enhancement in breadth of viral neutralization with these heterologous IM/IN groups was observed in the 129S1 mice compared to the other two strains when measured against B.1.351, BA.1, and especially against BA.4/5 (**Figs. 6B-E, G-J**). For example in the 129S1 mice, IM mRNA;IN NE/IVT/S treatment resulted in a 1 log enhancement in B.1.351 and BA.1 neutralization compared to the IM mRNA prime/boost group. The mouse strain difference was most apparent when the difference in nAb induction efficiency against the antigenically most distant BA.4/5 variant was considered. In 129S1 mice, all vaccine groups that included a prime boost had detectable BA.4/5-specific microneutralization titers except for mice that received IM mRNA prime/boost in which half of the group failed to show detectable neutralization (**Fig. 6E**). In K18-hACE2 mice, the same vaccination regimens resulted in less efficient induction of BA.4/5-specific nAb titers, with only the group that was primed with IM mRNA followed by IN NE/IVT/S boost inducing significant nAb titers in all of the animals within the immunization group (**Fig. 6J**). Thus, while the general trends are the same amongst the three mouse strains, there are nuanced differences which highlight the importance of comparing responses in the context of different genetic backgrounds. Finally, while IM Advx/S prime/boost induced similar total binding S-specific IgG titers as IM mRNA prime/boost, IN NE/IVT prime/boost, and IM mRNA prime with IN NE/S or NE/IVT/S boost, the IM Advx/S group displayed lower viral neutralization titers overall compared to these treatment groups. For example, in K18-hACE2 mice, the majority of IM Advx/S prime/boost mice showed no detectable neutralization of BA.4/5, pointing to the reduced breath of the antibody response induced by this adjuvant (**Fig. 6J**).

**Figure 6.**
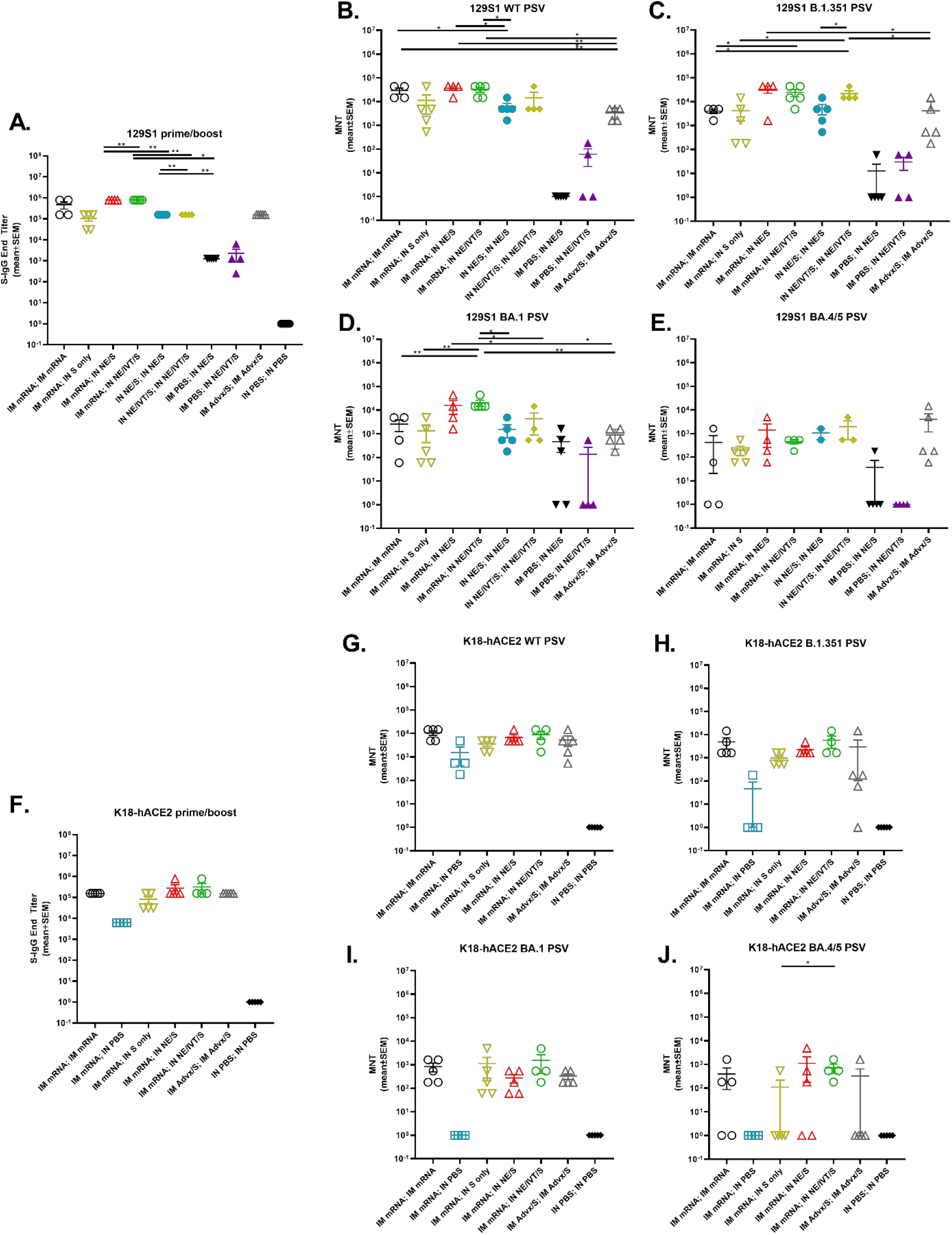
Heterologous IM/IN prime-boost immunization induces robust virus-neutralizing antibody titers in 129S1 and K18-hACE mice. 129S1 and K18-hACE2 mice were vaccinated twice 4-wks apart. Mice were primed either IM with 0.25μg of BNT162b2 mRNA or PBS, or IN with 15 μg full-length S protein with either NE or NE/IVT. Mice were then boosted 4 wks later IM with 0.25μg of BNT162b2 mRNA, or IN with 15 μg S with PBS, NE or NE/IVT as indicated. Groups receiving two immunizations with IM Advx/S or PBS were included for comparison. Serum **(A)** IgG titers against WT S-protein, and nAb titers against **(B)** WT, **(C)** B.1.351, **(D)** B.1.1.529 (BA.1), and **(E)** BA.4/5 variant PSVs were measured 2wks after the boost immunization (wk6). K18-hACE2 transgenic mice were similarly primed either IM with 0.25μg of BNT162b2 mRNA, or PBS, or IN with 15 μg S with either NE or NE/IVT. Mice were boosted 4 wks later IM with 0.25μg of BNT162b2 mRNA or PBS, or IN with 15 μg S with PBS, NE or NE/IVT as indicated. Groups receiving two immunizations with IM Advx/S or IN PBS were included for comparison. Serum **(F)** IgG titers against WT S-protein, and nAb titers against **(G)** WT, **(H)** B.1.351, **(I)** B.1.1.529 (BA.1), and **(J)** BA.4/5 variant PSVs were measured 2wks after the boost immunization (wk6). (n=4-5/grp;\**p<0.05*, \*\**p<0.01* by Mann-Whitney U test shown only for select groups-(full statistical analysis is shown in **Table S1**))

To evaluate protection against cross-variant viral challenge, we then infected 129S1 mice with 10^4^ PFU of B.1.351 (**Fig. 7A, B**) and K18-hACE2 mice with 10^4^ PFU of BA.5 (**Fig. 7C, D**) SARS-CoV-2 virus. Lungs as well as nasal turbinates (NTs) were harvested at 4 dpi for viral load determination by plaque assay. B.1.351 contains the N501Y mutation which allows it to replicate in WT 129S1 mice, causing up to 10% body weight loss.^28^ In contrast, while SARS-CoV-2 Omicron variants, including BA.5, can replicate in the respiratory tract of transgenic K18-hACE2 mice, infection is mainly characterized by the absence of overt morbidity.^29^ Nonetheless, it provides a useful model for evaluating breadth of immune protection with virus titer reduction as a surrogate of protection. In B.1.351 challenged 129S1 mice, IM mRNA prime/boost resulted in complete protection in the lungs (lower respiratory tract (LRT)) with the absence of replicating virus (**Fig. 7A**). However, the IM mRNA prime/boost vaccination failed to prevent viral replication in the nasal turbinates (upper respiratory tract (URT)) (**Fig. 7B**). In contrast, IM mRNA prime with IN adjuvanted S booster vaccination (NE/S, NE/IVT/S) demonstrated sterilizing immunity in both the LRT and URT, with complete absence of replicating virus in both the lungs and NTs of challenged mice. The IN NE/IVT/S prime/boost regimen also conferred sterilizing immunity in both the LRT and URT, highlighting the critical role of mucosal immunization in promoting URT protection. IN NE/S prime/boost immunization also conferred sterilizing immunity in the NTs, as well as significant protection in the lungs with most mice showing no viral replication in the lungs. However, 2/5 mice in this treatment group demonstrated modest breakthrough viral replication just above the limit of detection, supporting the advantage of the NE/IVT combined adjuvant. While IM Advx/S prime/boost offered a significant degree of protection as compared to the unvaccinated PBS control group, the Advx group showed the highest viral load in the lungs compared to all the other groups which received two immunizations. Furthermore, all mice in the Advx group demonstrated high viral titers in the NTs that were not significantly different from the unvaccinated control. These results further support the role of IN vaccination in promoting sterilizing immune responses in the URT. Notably, a single immunization with IN NE/S or NE/IVT/S did not confer significant protection in the lungs, yielding similar viral titers as the unvaccinated PBS group. However, a few animals in these groups did show a lack of viral replication in the NTs, while the others had similar titers as the unvaccinated control. Thus, the robust protective effects observed for the heterologous immunization groups is attributable to the synergistic effects of both the IM mRNA prime and NE-based IN pull components.

**Figure 7:**
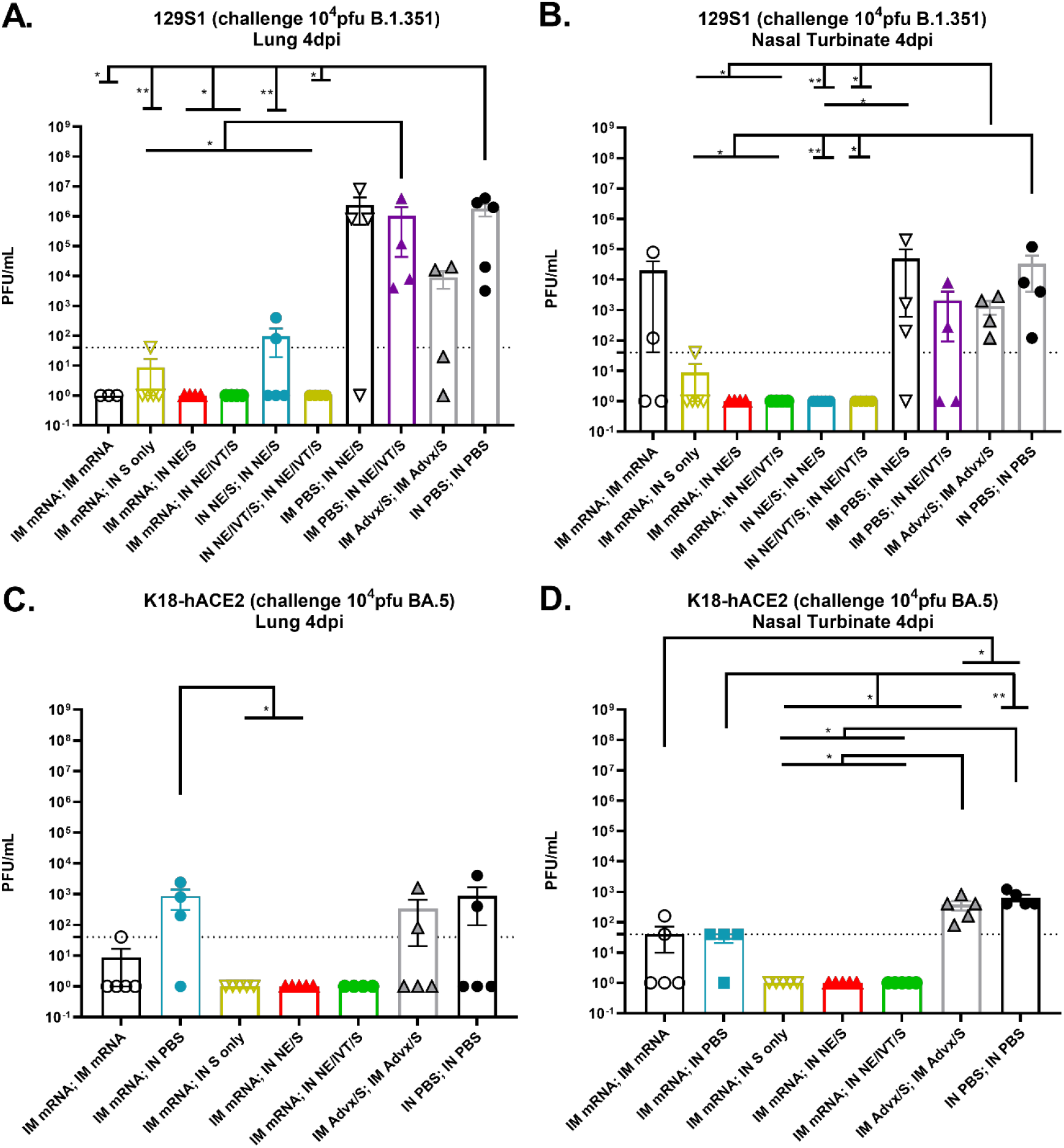
Heterologous IM/IN prime/pull and IN/IN immunization strategies provide sterilizing immunity upon heterologous challenge in both the upper and lower respiratory tracts in contrast to IM/IM immunization with BNT162b2 mRNA or Addavax/S. 129S1 mice were primed either IM with 0.25μg of BNT162b2 mRNA or PBS, or IN with 15 μg S with either NE or NE/IVT. Mice were boosted 4 wks later IM with 0.25μg of BNT162b2 mRNA, or IN with 15 μg S with PBS, NE or NE/IVT as indicated. Groups receiving two immunizations with IM Advx/S or IN PBS were included for comparison. 3 wks post-boost immunization, mice were challenged IN with 10^4^ pfu B.1.351, and viral titers were measured at 4 d.p.i. in the **(A)** lungs and **(B)** nasal turbinates. K18-hACE2 mice were primed IM with 0.25μg of BNT162b2 mRNA and boosted 4 wks later IM with 0.25μg of BNT162b2 mRNA, or IN with PBS alone, or 15 μg S with PBS, NE or NE/IVT as indicated. Groups receiving two immunizations with IM Advx/S or IN PBS were included for comparison. 3 wks post-boost immunization, mice were challenged IN with 10^4^ pfu BA.5, and viral titers were measured at 4 dpi in the **(A)** lungs and **(B)** nasal turbinates. (n=4-5/grp;\**p<0.05*, \*\**p<0.01* by Mann-Whitney U test)

Similar sterilizing immunity was observed for the IM mRNA prime, IN NE/S or IN NE/IVT/S boosted K18-hACE2 mice challenged with BA.5, with no viral replication detected in the lungs or NTs (**Fig. 7C, D**). Consistent with the results from the B.1.351 challenge in 129S1 mice, IM mRNA prime/boost also showed strong protection in the lungs after BA.5 challenge in vaccinated K18-hACE2 mice, as only 1/5 mice had detectable viral titers just above the detection limit. These mice also displayed higher levels of viral replication in the NTs compared to the IM mRNA prime, IN adjuvanted S groups showing low, but detectable viral titers in 2/5 mice. Similar to the B.1.351 challenged mice, the IM Advx/S prime/boost group also provided some degree of protection in the lungs of BA.5 challenged K18-hACE2 mice but had 2/5 mice with viral titers that were close those of unvaccinated controls. Moreover, no protection against BA.5 challenge in the NTs was observed in this group compared to unvaccinated controls. Notably, overall viral titers were lower for the BA.5 challenged K18-hACE2 mice even for the unimmunized control group as compared to B.1.351 challenged 129S1 mice, reflecting the poor infectivity of the Omicron variants in mouse models as others have previously observed^35,36^. Interestingly, IM mRNA prime, IN S only boost also provided sterilizing immunity to challenge in both the LRT and URT, for K18-hACE2 mice, as well as 129S1 mice, with only 1/5 129S1 mice showing viral titers at the detection limit (**Figs. 7A, B**). Thus, while the IM mRNA, IN adjuvanted S groups showed the most complete cross-variant sterilizing immune responses throughout the respiratory tract, such hybrid immunization approaches show a benefit in promoting protective mucosal immune responses even with unadjuvanted antigen alone delivered IN.

Host immune responses upon infection reflect disease course and pathogenesis, or lack thereof due to vaccine-mediated protection. Thus, we measured cytokine levels in lung homogenates at 4 d.p.i. for the vaccinated and challenged 129S1 and K18-hACE2 mice (**Figs. 8A, B**; individual cytokine data are provided in **Figs. S5, S6**). In 129S1 mice, pro-inflammatory innate cytokines/chemokines MIP-1α, MIP-1β, IP-10, MIP-2α, MCP-1, MCP-3, RANTES, GRO-α, IL-6 and TNFα were elevated in mice that showed breakthrough infection after receiving single vaccinations with either IN NE/S or IN NE/IVT/S, as well as with IM prime/boost with Advx/S or in the negative control group that received IN PBS twice (**Fig. 8A**). The T cell cytokines IFN-γ, IL-18, IL-22, and IL-10 were also increased in these groups, suggesting strong immune activation to control viral infection. Similarly, IM mRNA primed mice boosted IN with S alone or with NE/S had elevated innate cytokine/chemokine responses although with reduced adaptive immune cytokines. In contrast, 129S1 mice given homologous IM mRNA or IN NE/IVT/S prime/boost immunizations as well as mice given heterologous IM mRNA prime followed by IN NE/IVT/S boost had very low levels of both innate and adaptive chemokines/cytokines in the lungs post-challenge, consistent with the effective viral control observed in these groups. Furthermore, 129S1 mice that received IM Advx/S showed elevated T_H_2 cytokines IL-4, IL-5, IL-13 and eotaxin, which appeared higher than the unvaccinated group, reflecting the T_H_2 bias of the adjuvant. Unlike B.1.351 infection in 129S1 mice which resulted in strong host immune responses, BA.5 infection resulted in overall lower immune responses in K18-hACE2 mice, reflecting the better replication efficiency and pathogenesis of B.1.351 compared to BA.5 (**Figs. 8A, B, Figs. S5, S6)**. In K18-hACE2 mice, breakthrough infection in mice singly vaccinated with IM mRNA resulted in strongest induction of the innate pro-inflammatory cytokines/chemokines, MIP-1α, MIP-1β, IP-10, MCP-1, MCP-3, and IL-6 along with the cytokines TNF-α, IFN-γ, and IL-18. In contrast, similar to the 129S1 mice, groups primed with IM mRNA, boosted either IM with mRNA, or IN with NE/S or NE/IVT/S showed minimal induction of these inflammatory chemokines and cytokines, reflecting the effective viral control in these groups. Interestingly, IM mRNA prime followed by IN S alone showed a similar pro-inflammatory chemokine/cytokine profile as the single IM mRNA immunization group, albeit slightly reduced, highlighting the importance of the IN NE and NE/IVT adjuvants in mediating optimal protection. Overall, cytokine/chemokine profiles are elevated in groups for which replicating virus could be detected in lungs or nasal turbinates at 4 d.p.i., and chemokine/cytokine responses are skewed both by adjuvant type and vaccination routes.

**Figure 8:**
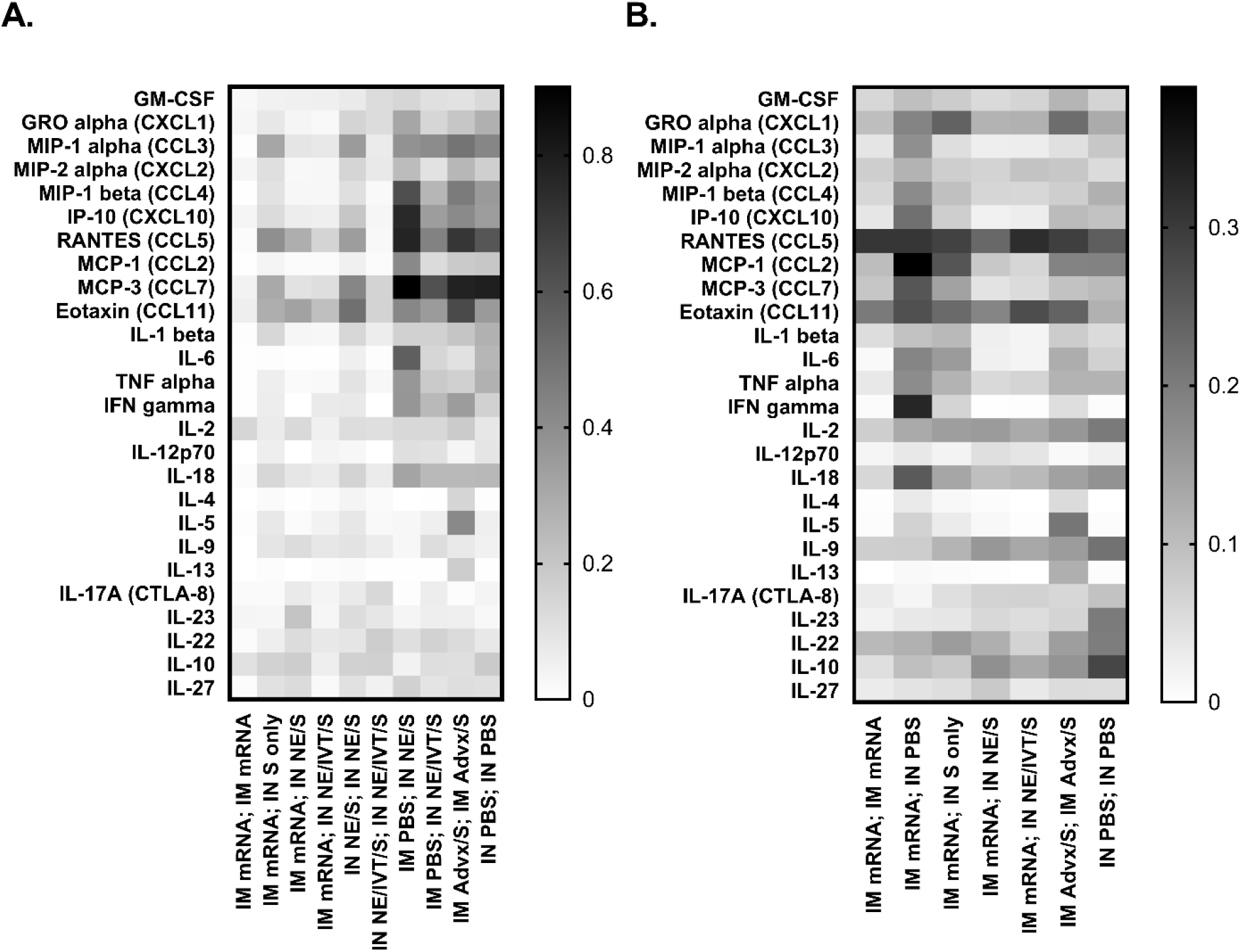
Cytokine/chemokine levels in lung homogenates from immunized mice post-challenge demonstrate different host response skewing depending on vaccination type and route. Cytokine and chemokine levels in lung homogenate measured by multiplex immunoassay from immunized (A) 129S1 mice in Figure 7 measured at 4 d.p.i. with 10^4^ pfu B.1.351, (B) K18-hACE2 mice in Figure 7 measured at 4 d.p.i. with 10^4^ pfu BA.5. Individual cytokine/chemokine levels were normalized to the cytokine/chemokine range and then normalized based on multiplication with log2 fold changes to normalize expression changes. Heatmap shows expression changes for the mean of each group. Individual cytokine levels and statistical analyses are provided in **Figures S5 and S6**.

## DISCUSSION

We have previously demonstrated the advantages of targeting multiple innate immune pathways for mucosal vaccination against SARS-CoV-2 utilizing a combined nanoemulsion/RIG-I agonist-based adjuvant containing subunit antigens including the full-length S protein, S1 subunit or RBD (REF). These vaccines were efficacious in inducing both circulating and mucosal antibodies and durable T cell responses which correlated with broad protection against infection with divergent SARS-CoV-2 variants even in the context of senescence.^15,16^ Currently licensed IM administered mRNA-based vaccines are efficient inducers of systemic nAbs and protective T cell responses but induce poor protective mucosal immune responses. However, evidence has suggested that “hybrid immunity” acquired through IM vaccination followed by natural infection provides more complete protection systemically and at mucosal sites.^30–32^ In an effort to emulate this hybrid immunity we assessed whether boosting via the IN route with NE/IVT-adjuvanted S-protein in IM mRNA-primed animals, could confer robust immunity through driving more optimal humoral and cellular immune responses, in *both* the periphery and at mucosal sites.

Humoral responses in serum were compared between various heterologous IM/IN and homologous IM/IM and IN/IN immunization regimens. In C57Bl/6 mice, priming with IM mRNA induced higher antigen-specific IgG than the IN NE or NE/IVT/S. After boost vaccination however, IM mRNA prime/boost, IN NE/IVT/S prime/boost and IM mRNA prime followed by heterologous boost with IN NE/S or NE/IVT/S resulted in equally robust S- and RBD-specific IgG titers. In contrast, minimal increase in antigen-specific IgG was observed in IM mRNA primed animals boosted IN with unadjuvanted S alone, highlighting the role of the NE adjuvants in driving an optimal boost effect for IN immunization. Similar effects were observed in mice of different genetic backgrounds. Furthermore, heterologous boosting with IN NE/S or NE/IVT/S induced robust cross-neutralizing antibody responses across WT, B.1.617.2, B.1.351, and B.1.1.529 variants which were enhanced compared to those induced by IM mRNA; IN S alone in IM mRNA primed C57Bl/6 and K18-hACE2 mice, confirming the effectiveness of the NE adjuvants in IN booster vaccines. The same relative patterns for nAbs were maintained in 129S1 mice, however, the IM mRNA; IN NE/IVT/S immunization showed clearer enhancement in viral cross-neutralizing antibodies compared to IM mRNA prime/boost. Such nuanced differences in the 129S1 mice may reflect the comparatively lower levels of IFNα production observed by others in C57Bl/6 and K18-hACE2 mice.^33,34^ Taken together, these results support the ability of the IN adjuvants to improve the breadth of the humoral immune response compared to homologous mRNA prime/boost regimens.

We have previously shown that the combination of NE and IVT results in a more T_H_1-polarized host immune response compared to the balanced T_H_1/T_H_2 response induced with NE alone, which is reflected in the antigen-specific IgG2b/IgG1 and IgG2c/IgG1 ratios in serum.^14^ Induction of IgG2c is significant due to its role in Fc-mediated effector functions including ADCC, ADCP, and complement activation which have shown to be key contributors to SARS-CoV-2 immunity.^35^ IN NE/IVT/S prime/boost induced higher IgG2b and IgG2c titers compared to IN NE/S prime/boost, confirming the T_H_1 skewing of the RIG-I agonist. However, IM mRNA prime followed by IM mRNA boost or IN boost with NE/S or NE/IVT/S resulted in similar IgG2b and significantly greater IgG2c as compared to prime/boost vaccination with IN NE/IVT/S, suggesting that priming with IM mRNA can enhance the T_H_1 polarization effects induced by IN NE/IVT. Indeed, T cell antigen-recall assessment in splenocytes and cLN isolates from vaccinated mice revealed dramatic enhancement of T_H_1 cytokine production by the heterologous IM mRNA; IN NE/IVT/S vaccination regimen compared to IM mRNA prime/boost, with high levels of secreted antigen-specific IFN-γ, IL-2, TNF-α, and IP-10 observed in response to S protein stimulation of splenocytes, with particularly high levels of these cytokines in the local mucosal draining lymph nodes. For example, IM mRNA; IN NE/IVT/S resulted in a nearly 20-fold enhancement in antigen-specific IFN-γ production in the cLN compared to IM mRNA prime/boost, demonstrating the crosstalk between the immune responses primed in the periphery by the mRNA and those triggered within the mucosa by the NE/IVT adjuvanted boost. Notably, IN boosting of IM mRNA primed T cell responses with NE/S or NE/IVT/S induced significantly higher percentages of S-specific polyfunctional IFN-γ^+^IL-2^+^TNF-α^+^CD4^+^ T cells in the cLN compared to homologous IM mRNA, IN NE/S, or IN NE/IVT/S prime/boost regimens. This is significant, as polyfunctional IFN-γ^+^IL-2^+^TNF-α^+^T cells have been shown to be a strong predictive indicator of potent antiviral T-cell responses. CD4^+^ T cells in secondary lymphoid tissues are required for class switching to result in production of IgG and IgA. Therefore, the optimal induction of CD4^+^ T cell responses in cLN is expected to promote class switching events in mucosal associated lymphoid tissues like the cLN, which correlates with the higher levels of IgA observed in heterologously boosted groups.

Interestingly, while all three IM mRNA prime, IN boost groups (S alone, NE/S, NE/IVT/S) induced higher frequencies of polyfunctional CD4^+^ T cells in the lung compared to IM mRNA prime/boost, the IN S alone boost did not significantly enhance these cells within the cLN, which required the NE or NE/IVT adjuvants for optimal enhancement. These results highlight the enhanced mucosal cellular responses induced by this mucosal booster strategy and underscore the role of the NE adjuvants in potentiating these responses. While homologous IM mRNA prime/boost induced significant levels of T_H_2 cytokines, IL-5 and IL-13 (mainly driven by some high responders), heterologous IN boosting with NE/S or NE/IVT/S did not enhance these responses, and reduced these responses, consistent with the strongly T_H_1-polarizing effects of the NE/IVT adjuvant. Finally, NE/IVT is a strong inducer of T_H_17 responses in the context of IL-10 production. T_H_17 cells in the context of IL-10 have been shown to be critical in promoting high and sustained levels of IgA production at mucosal sites-particularly the lung, and in the establishment of resident memory T cells.^25^ Induction of IL-17A is exclusive to the IN route of NE or NE/IVT administration and is a key component of NE/IVT mediated immunity. Interestingly, boosting IM mRNA primed mice with IN NE/IVT/S resulted equivalent levels of IL-17A production in the cLN as two immunizations with IN NE/IVT/S, which was higher than that of mice given only one IN NE/IVT/S immunization. Accordingly, we found that the IN NE/S or NE/IVT/S boost of IM mRNA primed mice resulted in increased frequencies of IL-17A-secreting CD4^+^T cells in the lung. These findings highlight the ability to use parenteral prime immunizations to set the stage for induction of more robust local mucosal responses upon mucosal pull immunization.

Consistent with IL-17A production, when induction of mucosal S-specific sIgA in BALF was considered, we found that IN immunization, either as a homologous prime/boost or as part of a heterologous IM prime/IN boost regimen, was required to obtain detectable S-specific IgA,. IM mRNA prime/boost did not induce detectable S-specific IgA, while IN NE/S and IN NE/IVT/S prime/boost resulted in robust sIgA responses. Interestingly, while single IN NE/S or IN NE/IVT/S immunizations induced only low levels of sIgA, equivalently high antigen-specific IgA was induced upon boosting IM mRNA primed animals with IN NE/IVT/S to similar levels as those induced by two immunizations with IN NE/IVT/S. These results further underscore the potential of the NE/IVT adjuvanted pull approach to drive improved mucosal immune responses initially primed by IM mRNA vaccination. It is well described that IgG antibodies can exudate from the blood into the lung and indeed, groups with highest serum ELISA S-binding titers also had highest S-specific binding ELISA titers in BALF. This suggests that deep lung antibody-mediated protection during infection may also be provided by IgG and not only sIgA, although aside from their shared ability to mediate viral neutralization, they can provide protection via different mechanisms. For example, IgA-mediated protection can involve removal of immune complexes through immune exclusion, as well as triggering FcαR1-mediated immune mechanisms like respiratory burst from neutrophils.^36^ IgG-mediated protection can also be occur through activation of ADCC and ADCP through Fcγ receptor-mediated mechanisms.^37^ Thus, protection in the respiratory tract provided by IgA and IgG likely occurs through a combination of mechanisms including direct virus neutralization, opsonization and the activation of Fc receptor dependent mechanisms.

As the N501Y mutation of B.1.351 allows it to infect WT mice, we used this variant to challenge 129S1 mice for evaluation of the impact of different vaccination strategies on virus replication and virus-host responses. Sterilizing immunity in the lungs was only observed for IM mRNA prime/boost mice and IM mRNA primed mice receiving a boost immunization through the IN route. These results demonstrate that induction of potent antibody responses through IM, IN or a combination of both can result in sterilizing immunity in the lungs. Prime/boost with IM Advx/S did not impart sterilizing immunity in the lung, despite having similar nAb titers against the B.1.351 variant as the IM mRNA; IN S group and similar serum S-specific serum IgG titers against the ancestral S protein as groups demonstrating sterilizing immunity. Thus, additional components besides serum nAbs are key contributors to sterilizing immunity in the LRT. When viral replication in the URT (nasal turbinates) was examined, we found that a mucosal boost vaccination was required to provide sterilizing immunity. Despite full control of viral replication in the LRT for the IM mRNA prime/boost group, viral replication was still detected in the URT. Furthermore, replicating virus was detected in the URT of all mice vaccinated with IM Advx/S prime/boost. Full control of viral replication in the URT thus correlated with induction of sIgA and local cellular responses by the mucosal booster strategies.

We observed that host-immune responses to infection are influenced by the type of adjuvanted vaccination, even if sterilizing immunity was observed. Heterologous boosting of IM mRNA primed mice IN with NE/S or NE/IVT/S in 129S1 mice markedly suppressed induction of major inflammatory markers associated with severe disease upon challenge with B.1.351, particularly in the IN NE/IVT/S boosted group. These findings confirm the protection afforded by these vaccination strategies. Similarly, homologous IM mRNA and IN NE/IVT/S prime/boost groups showed a similar cytokine/chemokine profile post-infection in the lung, with effective prevention of virally induced inflammatory responses typically associated with more severe disease^33^. Boosting IM mRNA primed mice with IN/S alone or homologous IN prime/boosting with NE/S also reduced inflammatory responses to infection compared to singly vaccinated animals and PBS control mice, however, in accordance with the incomplete protection observed in these groups as assessed by viral titers, this suppression was only partial. Increases in cytokines associated with type II polarization (IL-4, IL-5 and IL-13) in lungs post-infection were seen in mice that received IM Advx-adjuvanted vaccine. Promotion of T_H_2-driven vaccine responses appear to translate to type II host immune responses in the lungs of infected mice. This is an observation that we also have reported recently for mouse experiments performed with Advx-adjuvanted licensed influenza vaccines and influenza virus challenge.^38^

There is currently no optimal animal model for studying infection with Omicron SARS-CoV-2 variants, as they give attenuated pathology even in transgenic K18-hACE2 mice.^29^ Nevertheless, the K18-hACE2 infection model provides a good representation of mild Omicron SARS-CoV-2 infection for testing vaccine effectiveness when virus titers are considered in the URT and LRT. Similar to observations with B.1.351 infection in 129S1 mice, a mucosal boost vaccination of IM mRNA primed animals was required to obtain sterilizing immunity in both the URT and LRT in BA.5-challenged K18-hACE2 mice. In contrast, IM mRNA given once or as a prime/boost as well as IM Advx/S prime/boost were insufficient. In the K18-hACE2 model, mice that received only a single IM mRNA immunization showed a typical macrophage inflammatory profile upon BA.5 infection, whereas groups receiving heterologous IM mRNA prime, IN NE/S or NE/IVT/S boost effectively prevented induction of this inflammatory profile, consistent with the sterilizing immunity offered by these groups. If vaccination can prevent replication of virus in the nasal turbinates, this is seen as a first step in efficient interference with virus transmission. Taken together, these results strongly demonstrate that heterolgous IN boost vaccination is able to induce optimal mucosal immunity that effectively controls viral replication in the URT. Our ongoing experiments in hamster transmission models will be important for validating this prime/pull vaccination strategy also blocks viral transmission.

The ability to induce cross-variant sterilizing immunity within the respiratory tract is critical both for limiting viral transmission and disease progression, particularly as new SARS-CoV-2 variants continue to emerge. We herein demonstrated that an IM mRNA-based prime followed by a mucosal pull approach with a rationally designed adjuvant and recombinant protein is an effective strategy for inducing robust, tailored, and cross-sterilizing antibody and T cell responses both systemically and locally within the mucosa of the upper and lower respiratory tracts. Thus, as annual booster vaccinations continue to be required for COVID-19, our work underscores the value of a mucosal boost vaccination and highlights the promising potential of the IN NE/IVT adjuvant for inducing more complete hybrid immune responses in previously vaccinated subjects.

## MATERIALS AND METHODS

### Adjuvants and antigen

NE was produced by emulsifying cetylpyridinium chloride (CPC) and Tween 80 at a 1:6 (w/w) ratio, with ethanol (200 proof), super refined soybean oil (Croda) and water using a high-speed homogenizer as previously described.^40,41^ The sequence and synthesis of IVT DI RNA has previously been described in detail.^17^ Briefly, IVT DI was *in vitro* transcribed using a HiScribe T7 High Yield RNA synthesis kit (New England Biolabs) followed by DNAse I clean-up with a TURBO DNA-free kit (Thermo-Fisher), and purification with an RNeasy purification kit (Qiagen). Recombinant WT SARS-CoV-2 full-length S protein and RBD (aa319-545) (derived from Wuhan-Hu-1) with C-terminal His tags were produced in Expi293F or ExpiCHO cells, respectively, and purified by the University of Michigan Center for Structural Biology as previously described.^42^ Addavax (MF59 similar) was obtained from Invivogen, and the BNT162b2 mRNA vaccine (Pfizer) was obtained through the NIH SAVE program.

### Cell lines

Vero E6 cells (ATCC) were maintained in DMEM supplemented with 10% heat inactivated fetal bovine serum (HI FBS) and 1X non-essential amino acids (NEAA). HEK293T cells expressing hACE2 (293T-hACE2) were obtained from BEI resources and maintained in HEK293T medium: (DMEM with 4 mM L-glutamine, 4500 mg/L L-glucose, 1 mM sodium pyruvate and 1500 mg/L sodium bicarbonate, 10% HI FBS and 100 IU penicillin, and 100 μg/mL streptomycin).

### Viruses

SARS-CoV-2 clinical isolate USA-WA1/2020 (BEI resources; NR-52281), and B.1.351 and BA.5 variant viruses were propagated in Vero E6 cells or Vero-TMPRSS2. All viral stocks were verified by deep sequencing. All work with authentic SARS-CoV-2 viruses were performed in certified BSL3 or ABSL3 facilities in accordance with institutional safety and biosecurity procedures.

### Lentivirus pseudotyped virus

Generation of pseudotyped lentiviruses (PSVs) expressing the SARS-CoV-2 S proteins from WT, B.1.351, B.1.617.2, and B.1.1.529 (BA.1), and BA.4/5 variants harboring GFP and luciferase reporter genes was performed as previously described for the WT PSV.^43^ Plasmids carrying the full-length SARS-CoV-2 spike protein from each variant containing a C-terminal 19 amino acid deletion to remove the ER retention signal were used for pseudotyping (Invivogen). Viral titers (TU/mL) across variants were determined by measuring PSV transduction of GFP in 293T-hACE2 cells.

### Animals

All animal procedures were approved by the Institutional Animal Care and Use Committees at the University of Michigan and Icahn School of Medicine at Mount Sinai and were carried out in accordance with these guidelines. 6-8-wk-old female C57Bl/6, 129S1 (Jackson Laboratory), or K18-hACE2 mice (bred in-house) were housed in specific pathogen-free conditions. Mice were acclimated for 2 wks prior to initiation of each study. For challenge studies, mice were transferred to ABSL3 facilities 1 wk prior to viral challenge.

### Immunization

Mice were anesthetized using isoflurane in a IMPAC6 precision vaporizer. For IN immunization, mice were given 12 μL (6 μL/nare) of each vaccine formulation, and for IM immunization, the vaccine was delivered in a 50 μL volume. Each group received prime and boost immunizations at a 4-wk interval. For C57Bl/6 mice, mice were primed IM with 0.25 μg BNT162b2 mRNA (Pfizer/BioNTech). Mice were boosted either through the IM route with the same dose of mRNA, or through the IN route with PBS or 15 μg of WT S protein in PBS, 20 % NE (w/v) (NE/S) or 20% NE with 0.5 μg IVT DI (NE/IVT/S). Immune responses were compared to mice given homologous prime/boost immunizations with IN NE/S, IN NE/IVT/S, or IM 50% Addavax/S with the same amount of S protein. Comparison groups primed with IM PBS and boosted with IN NE/S or NE/IVT/S were also included. Select immunization regimens using the same adjuvant/antigen doses were chosen for evaluation in 129S1 and K18-hACE2 mice.

Serum was obtained by saphenous vein bleeding 2 and 4 wks after the prime, and by cardiac puncture at the end of the experiment at wk 8. Bronchial alveolar lavage fluid (BALF) was obtained by lung lavage with 0.8 mL PBS containing protease inhibitors at wk 8.

### ELISA

Immunograde 96-well ELISA plates (Midsci) were coated with 100 ng S protein or RBD in 50 μL PBS/well overnight at 4°C, and then blocked in 5% non-fat dry milk/PBS for 1 h at 37°C. Sera (or BAL) from immunized mice were serially diluted in PBSB (PBS/0.1% BSA). Blocking buffer was removed, and serum dilutions were added and incubated for 2 h at 37°C followed by overnight incubation at 4°C. Plates were washed with PBST (0.05% Tween20), and alkaline phosphatase conjugated secondary antibodies diluted in PBSB were added (goat-anti-mouse IgG, IgG1, IgG2b, IgG2c, (IgA for BAL) Jackson Immuno Research Laboratories), and incubated 1h at 37°C. Plates were washed with PBST, and developed by incubation with p-nitrophenyl phosphate (pNPP) substrate in diethanolamine (ThermoFisher) at RT. Absorbance was measured at 405 nm, and titers were determined using a cutoff defined by the sum of the average absorbance at the lowest dilution of naïve serum and two times the standard deviation.

### Pseudovirus microneutralization (MNT) assays

Pseudovirus (PSV) MNT assays were performed as previously described.^16^ Briefly, 1.25×10^4^ HEK293T-hACE2 cells/well were seeded overnight on 96-well tissue culture plates. Sera from immunized mice were serially diluted by a factor of three, starting at a dilution of 1:30 in HEK293T medium. 50 μL of diluted sera was added to 50 μL of PSVs (40,000 TU/mL) and incubated for 1h at 37°C. The PSV titer used across variant PSVs was selected based on the titer of WT PSV giving >100,000 RLUs above background. The virus/serum mixture was then added to the cells and incubated for 3 d at 37℃. Infection media was removed, and luminescence was measured by addition of 25 μL BrightGlo in 25 μL PBS. Neutralization titers were determined as the dilution at which the luminescence remained below the luminescence of the (virus only control-uninfected control)/2. Samples with undetectable neutralization were designated as having a titer of 10^0^. Neutralization assays with these PSVs have been demonstrated by us and numerous groups to be representative of authentic virus neutralization assays.^15,44^

### Tissue isolation and single cell suspension preparation

Two weeks post-boost (wk8), the left lung lobe was harvested, and single-cell suspensions were prepared by mincing with surgical scissors, and adding minced tissue to 3 mL of digestion media (RPMI, 10% HI FBS, 2 mM L-glutamine, 1% NEAA, 1 mM sodium pyruvate, 50 μM 2-mercaptoethanol, 100 IU penicillin, 100 μg/mL streptomycin with 1 mg/mL collagenase A (Roche), 20 U/mL of DNAse-I (Roche)) followed by incubation at 37°C for 1h with shaking. Tissue dissociation was continued by passages through an 18 gauge needle and filtering through a cell strainer. Cells were incubated in ACK lysis buffer for 5m at RT and washed with PBS. Methods for splenocyte and cLN lymphocyte preparation have previously been described^16^. All cells were resuspended in T cell media (DMEM, 5% HI FBS, 2 mM L-glutamine, 1% NEAA, 1 mM sodium pyruvate, 10 mM MOPS, 50 μM 2-mercaptoethanol, 100 IU penicillin, 100 μg/mL streptomycin) for further downstream analysis.

### Antigen Recall Response

T cell antigen recall response was assessed by cytokine release in cell isolates from the spleen and cLN of immunized mice 2 wks post-boost (wk 8). For antigen recall, isolated cells were plated at 8×10^5^cells/well and stimulated with 5 μg (25 μg/mL) S protein (WT) in T cell media for 72h at 37°C. Secreted cytokines (IFN-γ, IL-2, IP-10, IL-5, IL-6, IL-13, IL-10, IL-17A, and TNF-α) were measured relative to unstimulated cells in supernatants using a Milliplex MAP Magnetic Mouse Cytokine/Chemokine multiplex immunoassay (EMD Millipore).

### Flow Cytometry

For CD4^+^ intracellular cytokine analysis, 1×10^6^ cells isolated from the lungs, spleen, or cLNs were stimulated with 25 μg/mL S protein (WT) in T cell media for 24 h at 37°C, with brefeldin A (Biolegend) added during the last 6 h. Cells were stained with LIVE/DEAD Fixable Yellow Dead Cell Stain in PBS/2mM EDTA for 20 m at 4°C, washed, and then incubated with Fc block (Biolegend) in FACS buffer (PBS, 1% FBS, 2mM EDTA) for 10 m at 4°C prior to staining for surface T cell makers (CD3, CD4, CD8) for 30 m at 4°C. Cells were washed with FACS buffer, fixed in IC fixation buffer for 30 m at RT and permeabilized 30 m at 4°C before staining with antibodies for intracellular cytokines (**Table S2**) in permeabilization buffer for 30 m at 4°C. Cells were washed with permeabilization buffer and resuspended in PBS/2mM EDTA. FACS was performed on a Novocyte3000 flow cytometer and analyzed using FlowJo software.

### Viral Challenge

Immunized mice were challenged 3 weeks post-boost (wk 7). 129S1 mice were challenged IN with 10^4^ PFU B.1.351, and K18-hACE2 mice were challenged IN with 10^4^ PFU BA.5 delivered in 30 μL PBS. Mice were sacrificed 4 days post infection (d.p.i.). Lungs were harvested in 500 μL of PBS, and nasal turbinates were collected. Homogenates were prepared for virus titration by plaque assay as previously described, and cytokine profiling post-challenge was performed on homogenate using a Milliplex MAP Magnetic Mouse Cytokine/Chemokine multiplex immunoassay (EMD Millipore).^45^

### Statistical Analysis

Statistical analyses were performed with GraphPad Prism 9 (GraphPad Software). Comparisons were performed by Mann-Whitney U test or one-way ANOVA with Tukey post-hoc as indicated.

## Acknowledgements

SARS-CoV-2 work in the M.S. laboratory is supported by NIH/NIAID R01AI160706 and NIH/NIDDK R01DK130425. S.C.P has received support from the Korea Health Industry Development Institute (KHIDI) for research under the Biomedical Global Talent Nurturing Program (HI22C2101). Work in the P.T.W. laboratory is supported by NIH/NIAID R01AI160706 and NIH/NIAID R21AI176069. This study was also partly funded by CRIPT (Center for Research on Influenza Pathogenesis and Transmission), a NIH NIAID funded Center of Excellence for Influenza Research and Response (CEIRR, contract number 75N93021C00014), and by NIAID contract 75N93019C00046 to AG-S. We thank Richard Cadagan for technical support and Randy Albrecht for management and organization of the BSL3 facility. We thank Dr. Wendy Fonseca Aguilar for scientific input, the U of M vector core for producing the lentivirus pseudovirus and for providing technical input on assays, Joel Whitfield and the Rogel Cancer Center Immunology Core for multiplex assay support, as well as the U of M Center for Structural Biology for producing recombinant proteins. The CSB acknowledges support from the U-M Biosciences Initiative and the U-M Rogel Cancer Center. Finally, we also thank Daniel Flores, Marlene Espinoza, Jane Deng, and Ryan Camping for excellent administrative support.

## Author contributions

PTW, MS, GL, and MJW conceived of the studies and performed experiments, data analysis, and interpretation. DK, LAC, JJO, SP, PW, MF, JJL, and KWJ performed experiments and data analysis. PTW, MS, GL, MJW, and DK contributed to the writing of the manuscript, and JJO, AGS and JRB provided key scientific input.

