## Supplemental Info for "Enhanced mucosal B- and T-cell responses against SARS-CoV-2 after heterologous intramuscular mRNA prime/intranasal protein boost vaccination with a combination adjuvant"

**SUPPLEMENTAL INFORMATION:**

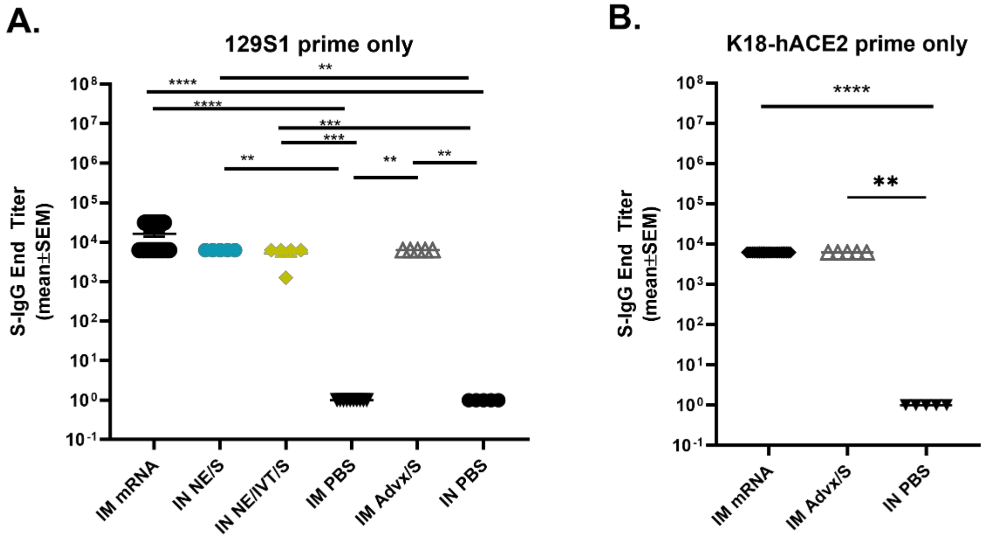

**Figure S1:** S-specific IgG induced 2 wks post-prime immunization in (A) 129S1 and (B) K18hACE2 mice immunized IM with 0.25µg of BNT162b2 mRNA or Advx with 15 µg S, or IN with 15 µg S with either NE or NE/IVT or PBS (n=5/grp; **p<0.05*, ***p<0.01*, ****p*<0.001, *****p*<0.0001 by Mann-Whitney U test).

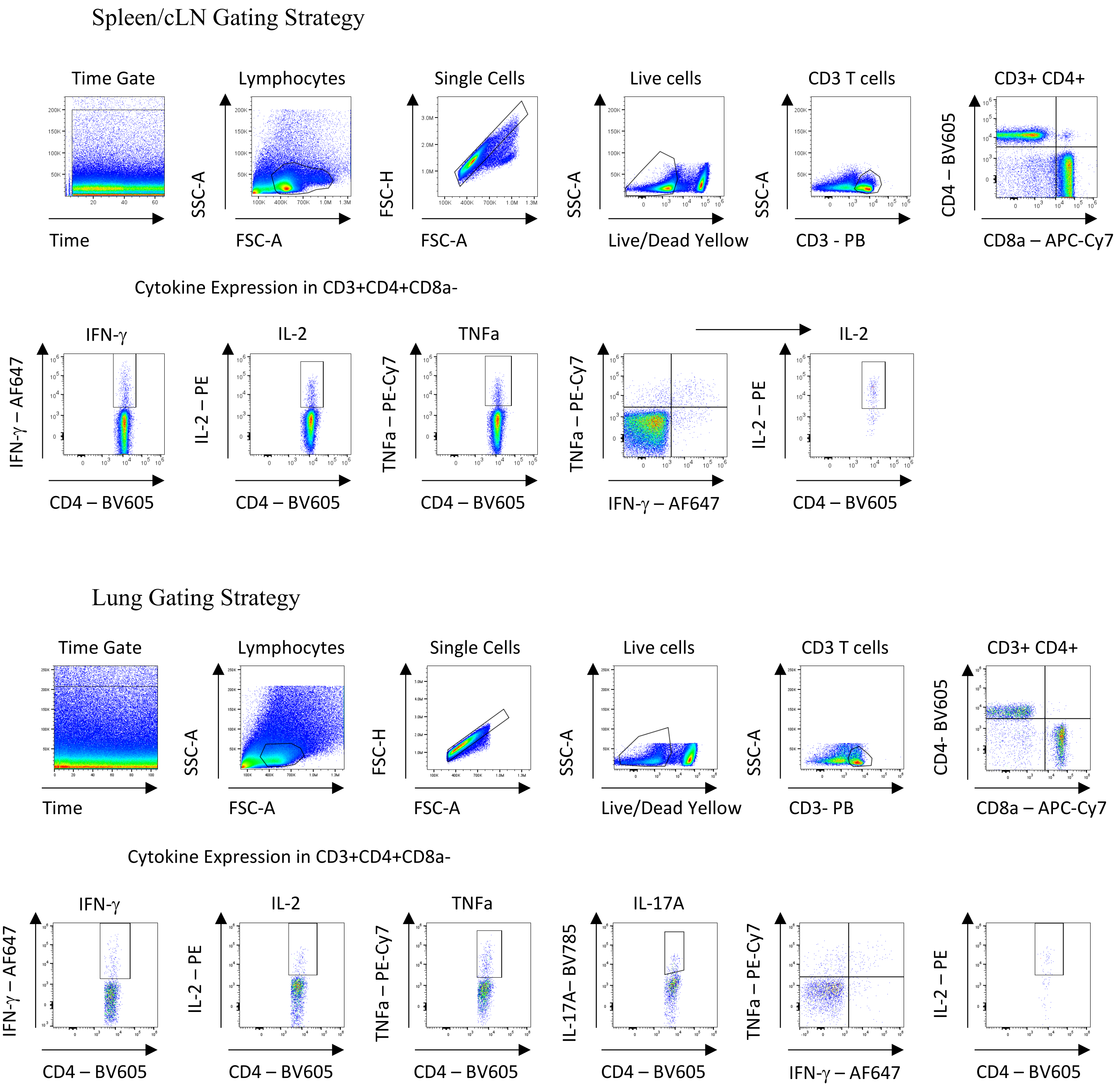

**Figure S2:** Gating strategy for ICS. Representative gating strategy for cytokine (IFN-γ, IL-2, TNFα, IL-17A) expression in CD4 T cells was evaluated in the (A) spleen, (B) cLN, and (C) lung after 24 hours of stimulation with 25μg/mL spike protein in the presence of Brefeldin A for the last 6 hours.

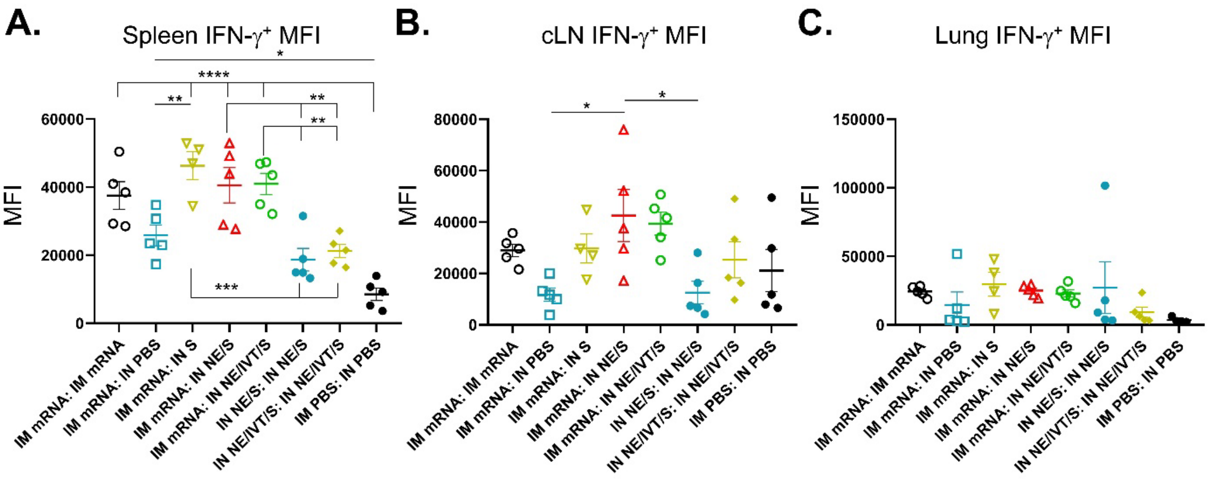

**Figure S3: MFI of IFN-γ expressing CD4 T cells.** Mean fluorescent intensity of IFN-γ expression in CD4 T cells in spleen, cLN and lung cells stimulated with 25μg/mL Spike protein for 24 hours. (n=4-5/grp; *$p<0.05$*, **$p<0.01$, ***$p<0.001$, ****$p<0.0001$ by One way ANOVA with Tukey post-hoc test.

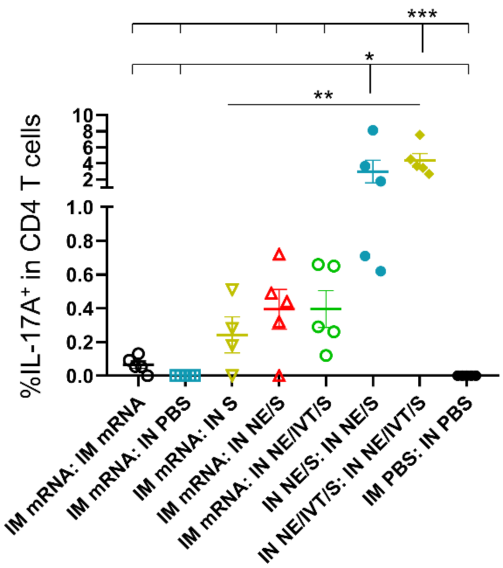

**Figure S4: IL-17A expression is highly induced in the lung by two doses of antigen with NE.** Single cell suspension were isolated from the lungs of mice immunized IM with 2µg BNT162b2 mRNA, or IN with 20µg of Spike protein in either PBS, NE, or NE/IVT. Cells were stimulated with 25µg/mL of S protein and IL-17A responses were quantified in CD4 T cells by intracellular cytokine staining. Data was analyzed by one-way ANOVA with Tukey post-hoc test. **p<0.05*, ***p<0.01, ***p<0.001.*

**Figure S5: Cytokine production in lung homogenates from 129S1 immunized mice post-challenge demonstrate different host response skewing depending on vaccination type and route.** Individual cytokine levels in lung homogenate (shown as heatmap in Figure 8) measured by multiplex immunoassay from immunized 129S1 mice in Figure 7 measured at 4 d.p.i. with $10^4$ pfu B.1.351. (A) IFN-γ, (B) IL-2, (C) TNF-α, (D) IL-12p70, (E) IP-10, (F) IL-4, (G) IL-5, (H) IL-13, (I) IL-6, (J) IL-17A, (K) IL-10, (L) IL-22, (M) IL-23, (N) IL-27, (O) IL-18, (P) IL-9, (Q) IL-1β, (R) MCP-1, (S) MCP-3, (T) MIP-1α, (U) MIP-1β, (V) MIP-2α, (W) RANTES, (X) GROα, (Y) GM-CSF, (Z) eotaxin (*n*=4-5/grp; *$p<0.05$, **$p<0.01$ by Mann-Whitney U test).

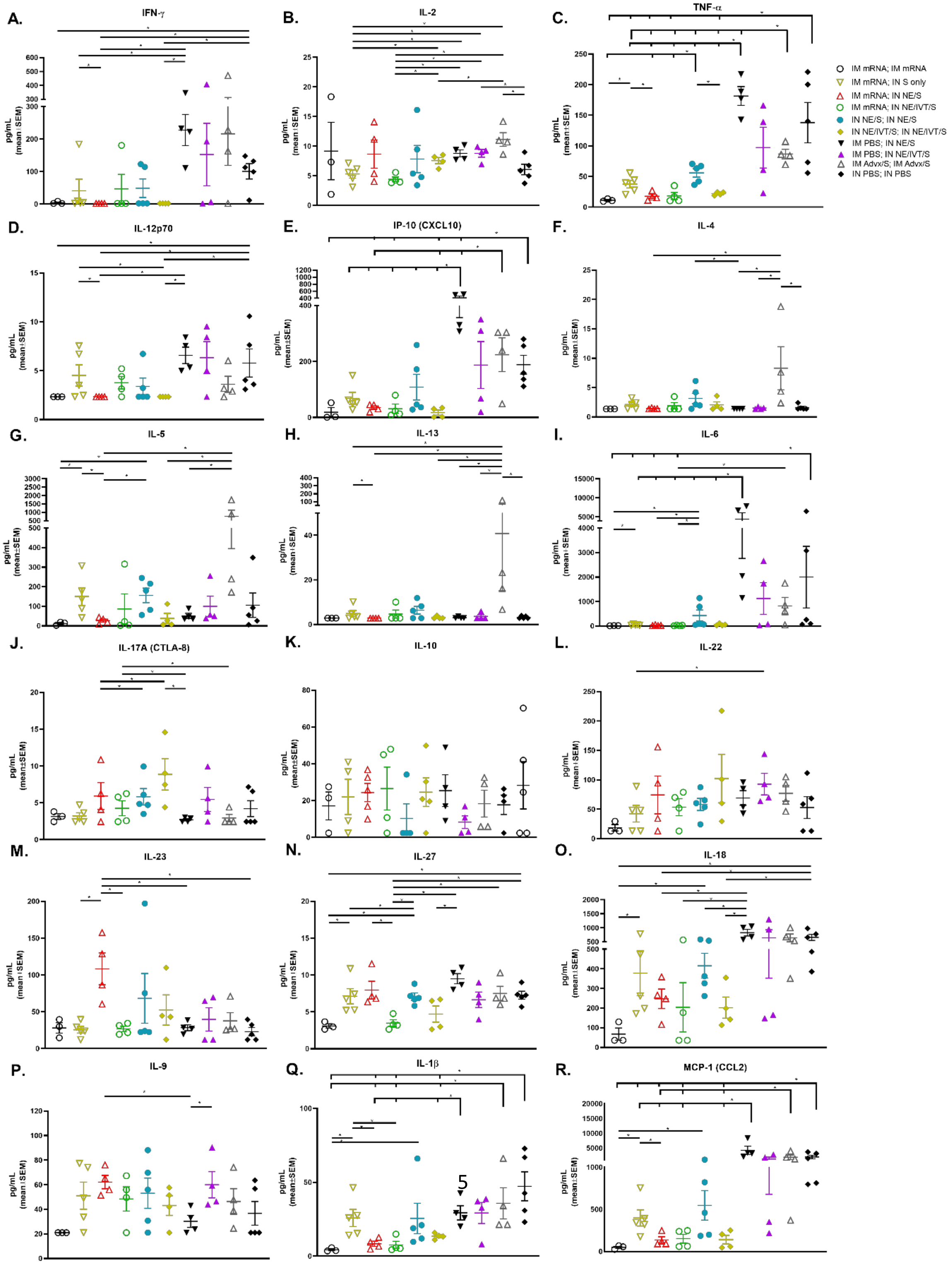

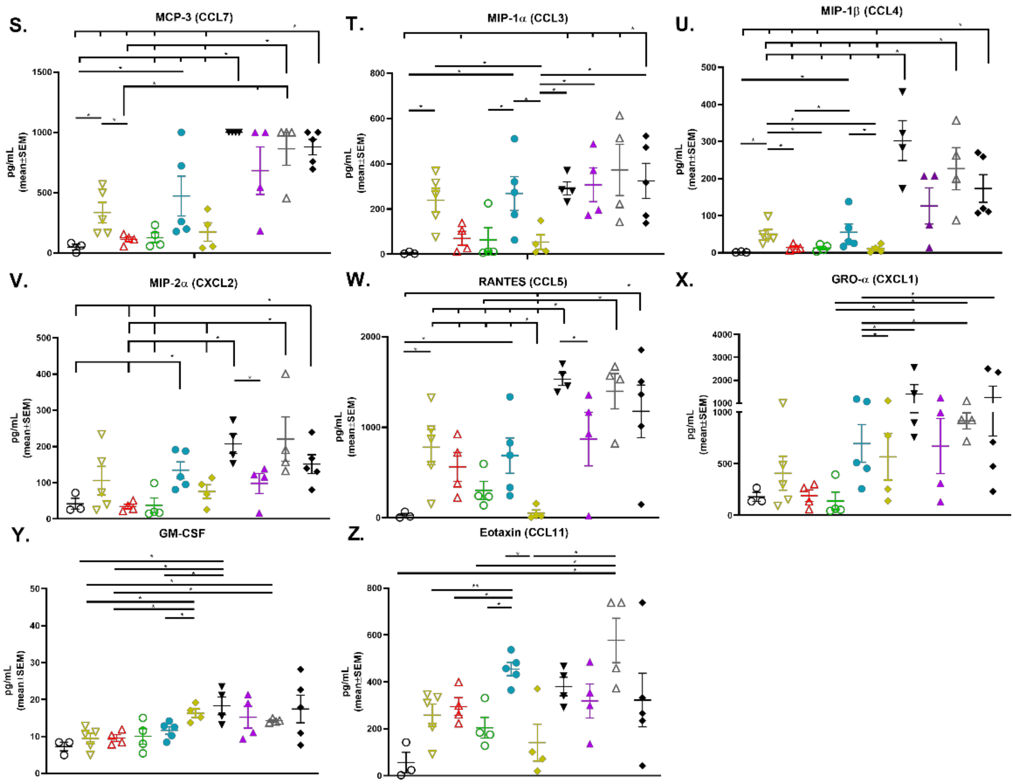
S.
MCP-3 (CCL7)
T.
MIP-1α (CCL3)
U.
MIP-1β (CCL4)
V.
MIP-2α (CXCL2)
W.
RANTES (CCL5)
X.
GRO-α (CXCL1)
Y.
GM-CSF
Z.
Eotaxin (CCL11)
pg/mL (mean±SEM)

**Figure S6: Cytokine production in lung homogenates from K18-hACE2 immunized mice post-challenge demonstrate different host response skewing depending on vaccination type and route.** Individual cytokine levels in lung homogenate (shown as heatmap in Figure 8) measured by multiplex immunoassay from immunized K18-hACE2 mice in Figure 7 measured at 4 d.p.i. with $10^4$ pfu BA.5. (A) IFN-γ, (B) IL-2, (C) TNF-α, (D) IL-12p70, (E) IP-10, (F) IL-4, (G) IL-5, (H) IL-13, (I) IL-6, (J) IL-17A, (K) IL-10, (L) IL-22, (M) IL-23, (N) IL-27, (O) IL-18, (P) IL-9, (Q) IL-1β, (R) MCP-1, (S) MCP-3, (T) MIP-1α, (U) MIP-1β, (V) MIP-2α, (W) RANTES, (X) GROα, (Y) GM-CSF, (Z) eotaxin (*n*=4-5/grp; **p<0.05*, ***p<0.01* by Mann-Whitney U test).

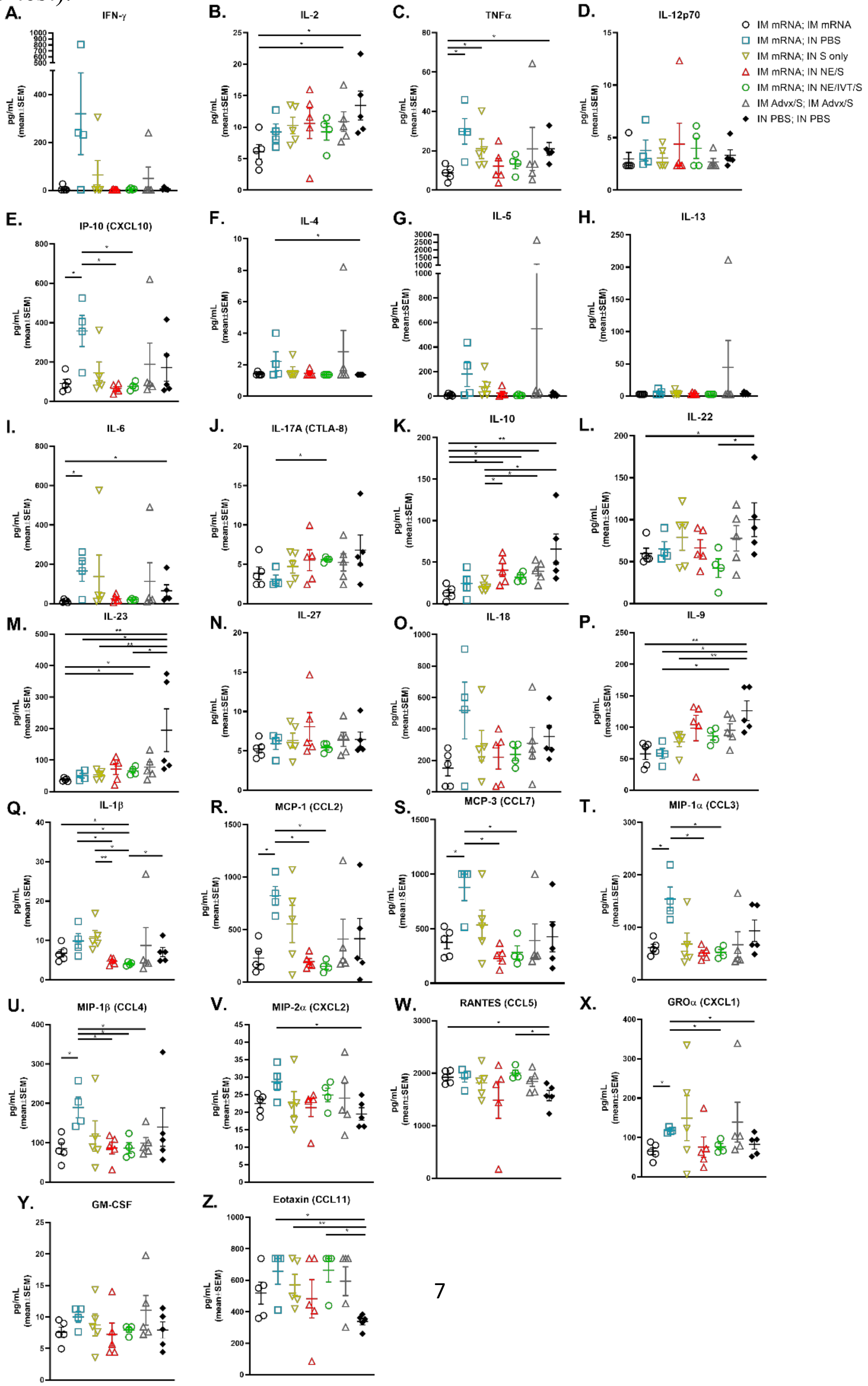

**TABLE S1:** Complete statistical analysis of data in Figures 1-5, and 6 (n=4-5/grp; *p<0.05, **p<0.01, ***p<0.001, ****p<0.0001 by Mann-Whitney U test or one way ANOVA with Tukey post-hoc test).

**Figure 1B:** C57Bl/6 wk 2 serum IgG against full-length WT S

| | IM mRNA | IN NE/S | IN NE/IVT/S | IM PBS |
|---|---|---|---|---|
| IM mRNA | | ** | **** | **** |
| IN NE/S | | | ns | *** |
| IN NE/IVT/S | | | | *** |
| IM PBS | | | | |

**Figure 1C:** C57Bl/6 wk 2 serum IgG against full-length WT RBD

| | IM mRNA | IN NE/S | IN NE/IVT/S | IM PBS |
|---|---|---|---|---|
| IM mRNA | | ns | ns | **** |
| IN NE/S | | | ns | *** |
| IN NE/IVT/S | | | | *** |
| IM PBS | | | | |

**Figure 1D:** C57Bl/6 wk 6 serum IgG against full-length WT S

| | IM mRNA; IM mRNA | IM mRNA; IN PBS | IM mRNA; IN S only | IM mRNA; IN NE/S | IM mRNA; IN NE/IVT/S | IN NE/S; IN NE/S | IN NE/IVT/S; IN NE/IVT/S | IM PBS; IN NE/S | IM PBS; IN NE/IVT/S |
|---|---|---|---|---|---|---|---|---|---|
| IM mRNA; IM mRNA | | ** | ** | ns | ns | * | ns | ** | ** |
| IM mRNA; IN PBS | | | ns | ** | ** | ** | ** | ns | ns |
| IM mRNA; IN S only | | | | * | ** | ns | * | ns | * |
| IM mRNA; IN NE/S | | | | | ns | * | ns | ** | ** |
| IM mRNA; IN NE/IVT/S | | | | | | * | ns | ** | ** |
| IN NE/S; IN NE/S | | | | | | | ns | ** | ** |
| IN NE/IVT/S; IN NE/IVT/S | | | | | | | | ** | ** |
| IM PBS; IN NE/S | | | | | | | | | ns |
| IM PBS; IN NE/IVT/S | | | | | | | | | |

**Figure 1E:** C57Bl/6 wk 6 serum IgG against full-length WT RBD

| | IM mRNA; IM mRNA | IM mRNA; IN PBS | IM mRNA; IN S only | IM mRNA; IN NE/S | IM mRNA; IN NE/IVT/S | IN NE/S; IN NE/S | IN NE/IVT/S; IN NE/IVT/S | IM PBS; IN NE/S | IM PBS; IN NE/IVT/S |
|---|---|---|---|---|---|---|---|---|---|
| IM mRNA; IM mRNA | | ** | ** | ns | ns | ns | ns | ** | ** |
| IM mRNA; IN PBS | | | ns | ** | ** | * | ** | ns | ns |
| IM mRNA; IN S only | | | | ** | ** | ns | ** | ns | ns |
| IM mRNA; IN NE/S | | | | | ns | ns | ns | ** | ** |
| IM mRNA; IN NE/IVT/S | | | | | | ns | ns | ** | ** |
| IN NE/S; IN NE/S | | | | | | | ns | * | ns |
| IN NE/IVT/S; IN NE/IVT/S | | | | | | | | ** | ** |
| IM PBS; IN NE/S | | | | | | | | | ns |
| IM PBS; IN NE/IVT/S | | | | | | | | | |

**Figure 1F:** C57Bl/6 wk 6 serum IgG1 against full-length WT S

| | IM mRNA; IM mRNA | IM mRNA; IN PBS | IM mRNA; IN S only | IM mRNA; IN NE/S | IM mRNA; IN NE/IVT/S | IN NE/S; IN NE/S | IN NE/IVT/S; IN NE/IVT/S | IM PBS; IN NE/S | IM PBS; IN NE/IVT/S |
|---|---|---|---|---|---|---|---|---|---|
| IM mRNA; IM mRNA | | ** | ** | ns | ns | ns | ns | ** | ** |
| IM mRNA; IN PBS | | | ns | * | * | * | ** | ns | ns |
| IM mRNA; IN S only | | | | ** | ** | ** | ** | ns | ns |
| IM mRNA; IN NE/S | | | | | ns | ns | ns | ** | ** |
| IM mRNA; IN NE/IVT/S | | | | | | ns | ns | ** | ** |
| IN NE/S; IN NE/S | | | | | | | ns | ** | ** |
| IN NE/IVT/S; IN NE/IVT/S | | | | | | | | ** | ** |
| IM PBS; IN NE/S | | | | | | | | | ns |
| IM PBS; IN NE/IVT/S | | | | | | | | | |

**Figure 1G:** C57Bl/6 wk 6 serum IgG2b against full-length WT S

| | IM mRNA; IM mRNA | IM mRNA; IN PBS | IM mRNA; IN S only | IM mRNA; IN NE/S | IM mRNA; IN NE/IVT/S | IN NE/S; IN NE/S | IN NE/IVT/S; IN NE/IVT/S | IM PBS; IN NE/S | IM PBS; IN NE/IVT/S |
|---|---|---|---|---|---|---|---|---|---|
| IM mRNA; IM mRNA | | ** | * | ns | ns | ns | ns | ** | ** |
| IM mRNA; IN PBS | | | ns | ** | ** | * | ** | ns | ns |
| IM mRNA; IN S only | | | | * | * | ns | * | * | * |
| IM mRNA; IN NE/S | | | | | ns | ns | ns | ** | ** |
| IM mRNA; IN NE/IVT/S | | | | | | ns | ns | ** | ** |
| IN NE/S; IN NE/S | | | | | | | * | ** | ** |
| IN NE/IVT/S; IN NE/IVT/S | | | | | | | | ** | ** |
| IM PBS; IN NE/S | | | | | | | | | ns |
| IM PBS; IN NE/IVT/S | | | | | | | | | |

**Figure 1H:** C57Bl/6 wk 6 serum IgG2c against full-length WT S

| | IM mRNA; IM mRNA | IM mRNA; IN PBS | IM mRNA; IN S only | IM mRNA; IN NE/S | IM mRNA; IN NE/IVT/S | IN NE/S; IN NE/S | IN NE/IVT/S; IN NE/IVT/S | IM PBS; IN NE/S | IM PBS; IN NE/IVT/S |
|---|---|---|---|---|---|---|---|---|---|
| IM mRNA; IM mRNA | | ** | ** | ns | ns | ** | ** | ** | ** |
| IM mRNA; IN PBS | | | ns | ** | ** | * | ns | ** | ** |
| IM mRNA; IN S only | | | | * | * | * | ns | ** | ** |
| IM mRNA; IN NE/S | | | | | ns | ** | * | ** | ** |
| IM mRNA; IN NE/IVT/S | | | | | | ** | * | ** | ** |
| IN NE/S; IN NE/S | | | | | | | ns | * | * |
| IN NE/IVT/S; IN NE/IVT/S | | | | | | | | * | * |
| IM PBS; IN NE/S | | | | | | | | | ns |
| IM PBS; IN NE/IVT/S | | | | | | | | | |

**Figure 1I:** C57Bl/6 wk 6 BAL IgA against full-length WT S

| | IM mRNA; IM mRNA | IM mRNA; IN PBS | IM mRNA; IN S only | IM mRNA; IN NE/S | IM mRNA; IN NE/IVT/S | IN NE/S; IN NE/S | IN NE/IVT/S; IN NE/IVT/S | IM PBS; IN NE/S | IM PBS; IN NE/IVT/S |
|---|---|---|---|---|---|---|---|---|---|
| IM mRNA; IM mRNA | | ns | ns | ** | * | * | * | ns | ns |
| IM mRNA; IN PBS | | | ns | ** | * | * | * | ns | ns |
| IM mRNA; IN S only | | | | ns | ns | ns | ns | ns | ns |
| IM mRNA; IN NE/S | | | | | ns | ns | ns | * | ** |
| IM mRNA; IN NE/IVT/S | | | | | | ns | ns | * | * |
| IN NE/S; IN NE/S | | | | | | | ns | * | * |
| IN NE/IVT/S; IN NE/IVT/S | | | | | | | | * | * |
| IM PBS; IN NE/S | | | | | | | | | ns |
| IM PBS; IN NE/IVT/S | | | | | | | | | |

**Figure 1J:** C57Bl/6 wk 6 BAL IgG against full-length WT S

| | IM mRNA; IM mRNA | IM mRNA; IN PBS | IM mRNA; IN S only | IM mRNA; IN NE/S | IM mRNA; IN NE/IVT/S | IN NE/S; IN NE/S | IN NE/IVT/S; IN NE/IVT/S | IM PBS; IN NE/S | IM PBS; IN NE/IVT/S |
|---|---|---|---|---|---|---|---|---|---|
| IM mRNA; IM mRNA | | ** | ** | ns | * | ** | ns | ** | ** |
| IM mRNA; IN PBS | | | ns | ** | ** | * | ** | ns | * |
| IM mRNA; IN S only | | | | * | * | ns | * | ns | * |
| IM mRNA; IN NE/S | | | | | ns | ns | ns | ** | ** |
| IM mRNA; IN NE/IVT/S | | | | | | ns | ns | ** | ** |
| IN NE/S; IN NE/S | | | | | | | ns | ** | ** |
| IN NE/IVT/S; IN NE/IVT/S | | | | | | | | ** | ** |
| IM PBS; IN NE/S | | | | | | | | | ns |
| IM PBS; IN NE/IVT/S | | | | | | | | | |

**Figure 2A**: wk6 serum WT PSV MNT

| | IM mRNA; IM mRNA | IM mRNA; IN PBS | IM mRNA; IN S only | IM mRNA; IN NE/S | IM mRNA; IN NE/IVT/S | IN NE/S; IN NE/S | IN NE/IVT/S; IN NE/IVT/S |
|---|---|---|---|---|---|---|---|
| IM mRNA; IM mRNA | | * | ** | ns | ns | * | ns |
| IM mRNA; IN PBS | | | ns | ** | * | * | * |
| IM mRNA; IN S only | | | | ** | * | * | ns |
| IM mRNA; IN NE/S | | | | | ns | ns | ns |
| IM mRNA; IN NE/IVT/S | | | | | | ns | ns |
| IN NE/S; IN NE/S | | | | | | | ns |
| IN NE/IVT/S; IN NE/IVT/S | | | | | | | |

**Figure 2B**: wk6 serum B.1.617.2 PSV MNT

| | IM mRNA; IM mRNA | IM mRNA; IN PBS | IM mRNA; IN S only | IM mRNA; IN NE/S | IM mRNA; IN NE/IVT/S | IN NE/S; IN NE/S | IN NE/IVT/S; IN NE/IVT/S |
|---|---|---|---|---|---|---|---|
| IM mRNA; IM mRNA | | ** | * | ns | ns | ns | ns |
| IM mRNA; IN PBS | | | ns | ** | * | * | ** |
| IM mRNA; IN S only | | | | * | ns | * | ** |
| IM mRNA; IN NE/S | | | | | ns | ns | ns |
| IM mRNA; IN NE/IVT/S | | | | | | ns | ns |
| IN NE/S; IN NE/S | | | | | | | ns |
| IN NE/IVT/S; IN NE/IVT/S | | | | | | | |

**Figure 2C**: wk6 serum B.1.351 PSV MNT

| | IM mRNA; IM mRNA | IM mRNA; IN PBS | IM mRNA; IN S only | IM mRNA; IN NE/S | IM mRNA; IN NE/IVT/S | IN NE/S; IN NE/S | IN NE/IVT/S; IN NE/IVT/S |
|---|---|---|---|---|---|---|---|
| IM mRNA; IM mRNA | | * | ** | ns | ns | ns | ns |
| IM mRNA; IN PBS | | | * | ns | ns | * | ns |
| IM mRNA; IN S only | | | | * | ns | ** | * |
| IM mRNA; IN NE/S | | | | | ns | ns | ns |
| IM mRNA; IN NE/IVT/S | | | | | | ns | ns |
| IN NE/S; IN NE/S | | | | | | | ns |
| IN NE/IVT/S; IN NE/IVT/S | | | | | | | |

**Figure 2D**: wk6 serum BA.1 PSV MNT

| | IM mRNA; IM mRNA | IM mRNA; IN PBS | IM mRNA; IN S only | IM mRNA; IN NE/S | IM mRNA; IN NE/IVT/S | IN NE/S; IN NE/S | IN NE/IVT/S; IN NE/IVT/S |
|---|---|---|---|---|---|---|---|
| IM mRNA; IM mRNA | | ns | ns | ns | ns | ns | ns |
| IM mRNA; IN PBS | | | ns | ns | ns | * | ns |
| IM mRNA; IN S only | | | | ns | ns | * | ns |
| IM mRNA; IN NE/S | | | | | ns | ns | ns |
| IM mRNA; IN NE/IVT/S | | | | | | ns | ns |
| IN NE/S; IN NE/S | | | | | | | ns |
| IN NE/IVT/S; IN NE/IVT/S | | | | | | | |

**Figure 3A**: SPL IFNg

| | IM mRNA; IM mRNA | IM mRNA; IN PBS | IM mRNA; IN S only | IM mRNA; IN NE/S | IM mRNA; IN NE/IVT/S | IN NE/S; IN NE/S | IN NE/IVT/S; IN NE/IVT/S | IM PBS; IN NE/S | IM PBS; IN NE/IVT/S |
|---|---|---|---|---|---|---|---|---|---|
| IM mRNA; IM mRNA | | ns | ns | ns | ns | * | ns | * | * |
| IM mRNA; IN PBS | | | ns | ns | ns | ns | ns | ns | * |
| IM mRNA; IN S only | | | | ns | * | ns | ns | ns | * |
| IM mRNA; IN NE/S | | | | | ns | ns | ns | ns | ns |
| IM mRNA; IN NE/IVT/S | | | | | | * | ns | ** | * |
| IN NE/S; IN NE/S | | | | | | | * | ns | * |
| IN NE/IVT/S; IN NE/IVT/S | | | | | | | | * | * |
| IM PBS; IN NE/S | | | | | | | | | ns |
| IM PBS; IN NE/IVT/S | | | | | | | | | |

**Figure 3B**: SPL IL2

| | IM mRNA; IM mRNA | IM mRNA; IN PBS | IM mRNA; IN S only | IM mRNA; IN NE/S | IM mRNA; IN NE/IVT/S | IN NE/S; IN NE/S | IN NE/IVT/S; IN NE/IVT/S | IM PBS; IN NE/S | IM PBS; IN NE/IVT/S |
|---|---|---|---|---|---|---|---|---|---|
| IM mRNA; IM mRNA | | ns | ns | ns | * | ns | ns | ns | * |
| IM mRNA; IN PBS | | | ns | ns | ns | ns | ns | ns | ns |
| IM mRNA; IN S only | | | | ns | ns | ns | ns | * | * |
| IM mRNA; IN NE/S | | | | | ns | * | ns | ** | * |
| IM mRNA; IN NE/IVT/S | | | | | | ** | ns | ** | * |
| IN NE/S; IN NE/S | | | | | | | ns | ns | ns |
| IN NE/IVT/S; IN NE/IVT/S | | | | | | | | * | * |
| IM PBS; IN NE/S | | | | | | | | | ns |
| IM PBS; IN NE/IVT/S | | | | | | | | | |

**Figure 3C**: SPL IP10

| | IM mRNA; IM mRNA | IM mRNA; IN PBS | IM mRNA; IN S only | IM mRNA; IN NE/S | IM mRNA; IN NE/IVT/S | IN NE/S; IN NE/S | IN NE/IVT/S; IN NE/IVT/S | IM PBS; IN NE/S | IM PBS; IN NE/IVT/S |
|---|---|---|---|---|---|---|---|---|---|
| IM mRNA; IM mRNA | | ns | ns | ns | ns | ns | ns | ns | ns |
| IM mRNA; IN PBS | | | ns | ns | ns | ns | ns | ns | ns |
| IM mRNA; IN S only | | | | ns | ns | ns | ns | ns | ns |
| IM mRNA; IN NE/S | | | | | ns | ns | ** | ns | ns |
| IM mRNA; IN NE/IVT/S | | | | | | ns | ns | ns | ns |
| IN NE/S; IN NE/S | | | | | | | ns | ns | ns |
| IN NE/IVT/S; IN NE/IVT/S | | | | | | | | ns | ns |
| IM PBS; IN NE/S | | | | | | | | | ns |
| IM PBS; IN NE/IVT/S | | | | | | | | | |

**Figure 3D**: SPL TNFa

| | IM mRNA; IM mRNA | IM mRNA; IN PBS | IM mRNA; IN S only | IM mRNA; IN NE/S | IM mRNA; IN NE/IVT/S | IN NE/S; IN NE/S | IN NE/IVT/S; IN NE/IVT/S | IM PBS; IN NE/S | IM PBS; IN NE/IVT/S |
|---|---|---|---|---|---|---|---|---|---|
| IM mRNA; IM mRNA | | ns | ns | ns | ns | ns | ns | ns | ns |
| IM mRNA; IN PBS | | | ns | ns | ns | ns | ns | ns | ns |
| IM mRNA; IN S only | | | | ns | ns | ns | ns | ns | ns |
| IM mRNA; IN NE/S | | | | | ns | ns | ns | ns | ns |
| IM mRNA; IN NE/IVT/S | | | | | | ns | ns | ns | * |
| IN NE/S; IN NE/S | | | | | | | ns | ns | * |
| IN NE/IVT/S; IN NE/IVT/S | | | | | | | | ns | * |
| IM PBS; IN NE/S | | | | | | | | | ns |
| IM PBS; IN NE/IVT/S | | | | | | | | | |

**Figure 3E**: SPL IL5

| | IM mRNA; IM mRNA | IM mRNA; IN PBS | IM mRNA; IN S only | IM mRNA; IN NE/S | IM mRNA; IN NE/IVT/S | IN NE/S; IN NE/S | IN NE/IVT/S; IN NE/IVT/S | IM PBS; IN NE/S | IM PBS; IN NE/IVT/S |
|---|---|---|---|---|---|---|---|---|---|
| IM mRNA; IM mRNA | | ns | ns | ns | ns | * | ** | ** | * |
| IM mRNA; IN PBS | | | ns | ns | ns | ns | ns | ** | * |
| IM mRNA; IN S only | | | | ns | ns | ns | ns | ** | * |
| IM mRNA; IN NE/S | | | | | ns | ns | ns | ** | * |
| IM mRNA; IN NE/IVT/S | | | | | | ns | ns | ** | * |
| IN NE/S; IN NE/S | | | | | | | ns | ** | * |
| IN NE/IVT/S; IN NE/IVT/S | | | | | | | | ** | * |
| IM PBS; IN NE/S | | | | | | | | | ns |
| IM PBS; IN NE/IVT/S | | | | | | | | | |

**Figure 3F**: SPL IL13

| | IM mRNA; IM mRNA | IM mRNA; IN PBS | IM mRNA; IN S only | IM mRNA; IN NE/S | IM mRNA; IN NE/IVT/S | IN NE/S; IN NE/S | IN NE/IVT/S; IN NE/IVT/S | IM PBS; IN NE/S | IM PBS; IN NE/IVT/S |
|---|---|---|---|---|---|---|---|---|---|
| IM mRNA; IM mRNA | | ns | ns | ns | ns | * | * | ** | * |
| IM mRNA; IN PBS | | | ns | ns | ns | ns | ns | ns | ns |
| IM mRNA; IN S only | | | | ns | ns | ns | ns | * | * |
| IM mRNA; IN NE/S | | | | | ns | ns | ns | * | * |
| IM mRNA; IN NE/IVT/S | | | | | | ns | ns | ** | * |
| IN NE/S; IN NE/S | | | | | | | ns | ns | ns |
| IN NE/IVT/S; IN NE/IVT/S | | | | | | | | * | ns |
| IM PBS; IN NE/S | | | | | | | | | ns |
| IM PBS; IN NE/IVT/S | | | | | | | | | |

**Figure 3G**: SPL IL6

| | IM mRNA; IM mRNA | IM mRNA; IN PBS | IM mRNA; IN S only | IM mRNA; IN NE/S | IM mRNA; IN NE/IVT/S | IN NE/S; IN NE/S | IN NE/IVT/S; IN NE/IVT/S | IM PBS; IN NE/S | IM PBS; IN NE/IVT/S |
|---|---|---|---|---|---|---|---|---|---|
| IM mRNA; IM mRNA | | ns | ns | ns | ** | ns | ns | ns | ns |
| IM mRNA; IN PBS | | | ns | ns | ** | ns | ns | ns | ns |
| IM mRNA; IN S only | | | | ns | ** | ns | ns | ns | ns |
| IM mRNA; IN NE/S | | | | | ns | * | ns | * | * |
| IM mRNA; IN NE/IVT/S | | | | | | ** | ** | ** | * |
| IN NE/S; IN NE/S | | | | | | | ns | ns | ns |
| IN NE/IVT/S; IN NE/IVT/S | | | | | | | | ns | ns |
| IM PBS; IN NE/S | | | | | | | | | ns |
| IM PBS; IN NE/IVT/S | | | | | | | | | |

**Figure 3H**: SPL IL17A

| | IM mRNA; IM mRNA | IM mRNA; IN PBS | IM mRNA; IN S only | IM mRNA; IN NE/S | IM mRNA; IN NE/IVT/S | IN NE/S; IN NE/S | IN NE/IVT/S; IN NE/IVT/S | IM PBS; IN NE/S | IM PBS; IN NE/IVT/S |
|---|---|---|---|---|---|---|---|---|---|
| IM mRNA; IM mRNA | | ns | * | ** | ** | * | ** | ** | * |
| IM mRNA; IN PBS | | | ns | ns | ** | ns | ** | ns | ns |
| IM mRNA; IN S only | | | | ns | ** | ns | ** | ns | ns |
| IM mRNA; IN NE/S | | | | | ns | ns | * | ns | ns |
| IM mRNA; IN NE/IVT/S | | | | | | ns | ns | ** | ns |
| IN NE/S; IN NE/S | | | | | | | ns | ns | ns |
| IN NE/IVT/S; IN NE/IVT/S | | | | | | | | ** | * |
| IM PBS; IN NE/S | | | | | | | | | ns |
| IM PBS; IN NE/IVT/S | | | | | | | | | |

**Figure 3I**: SPL IL10

| | IM mRNA; IM mRNA | IM mRNA; IN PBS | IM mRNA; IN S only | IM mRNA; IN NE/S | IM mRNA; IN NE/IVT/S | IN NE/S; IN NE/S | IN NE/IVT/S; IN NE/IVT/S | IM PBS; IN NE/S | IM PBS; IN NE/IVT/S |
|---|---|---|---|---|---|---|---|---|---|
| IM mRNA; IM mRNA | | ns | ns | ns | ns | ns | ns | ns | * |
| IM mRNA; IN PBS | | | ns | ns | ns | ns | ns | ns | ns |
| IM mRNA; IN S only | | | | ns | ns | ns | ns | ns | * |
| IM mRNA; IN NE/S | | | | | ns | ns | ns | ns | * |
| IM mRNA; IN NE/IVT/S | | | | | | ns | ns | ns | * |
| IN NE/S; IN NE/S | | | | | | | ns | ns | ns |
| IN NE/IVT/S; IN NE/IVT/S | | | | | | | | ns | * |
| IM PBS; IN NE/S | | | | | | | | | ns |
| IM PBS; IN NE/IVT/S | | | | | | | | | |

**Figure 4A:** cLN IFNg

| | IM mRNA; IM mRNA | IM mRNA; IN PBS | IM mRNA; IN S only | IM mRNA; IN NE/S | IM mRNA; IN NE/IVT/S | IN NE/S; IN NE/S | IN NE/IVT/S; IN NE/IVT/S | IM PBS; IN NE/S | IM PBS; IN NE/IVT/S |
|---|---|---|---|---|---|---|---|---|---|
| IM mRNA; IM mRNA | | ns | ns | ns | ** | ns | ns | ns | ns |
| IM mRNA; IN PBS | | | ns | * | ** | ns | * | ns | ns |
| IM mRNA; IN S only | | | | ns | ** | ns | ns | ns | ns |
| IM mRNA; IN NE/S | | | | | ns | ns | ns | ns | ns |
| IM mRNA; IN NE/IVT/S | | | | | | ** | ** | ** | ns |
| IN NE/S; IN NE/S | | | | | | | * | ns | ns |
| IN NE/IVT/S; IN NE/IVT/S | | | | | | | | ns | ns |
| IM PBS; IN NE/S | | | | | | | | | ns |
| IM PBS; IN NE/IVT/S | | | | | | | | | |

**Figure 4B:** cLN IL2

| | IM mRNA; IM mRNA | IM mRNA; IN PBS | IM mRNA; IN S only | IM mRNA; IN NE/S | IM mRNA; IN NE/IVT/S | IN NE/S; IN NE/S | IN NE/IVT/S; IN NE/IVT/S | IM PBS; IN NE/S | IM PBS; IN NE/IVT/S |
|---|---|---|---|---|---|---|---|---|---|
| IM mRNA; IM mRNA | | ns | ** | ** | ** | ** | ** | ** | * |
| IM mRNA; IN PBS | | | ns | * | * | ns | * | ns | ns |
| IM mRNA; IN S only | | | | ns | ns | ns | * | ns | ns |
| IM mRNA; IN NE/S | | | | | ns | ns | ns | * | * |
| IM mRNA; IN NE/IVT/S | | | | | | ns | ns | ns | * |
| IN NE/S; IN NE/S | | | | | | | ns | ns | ns |
| IN NE/IVT/S; IN NE/IVT/S | | | | | | | | * | * |
| IM PBS; IN NE/S | | | | | | | | | ns |
| IM PBS; IN NE/IVT/S | | | | | | | | | |

**Figure 4C:** cLN IP10

| | IM mRNA; IM mRNA | IM mRNA; IN PBS | IM mRNA; IN S only | IM mRNA; IN NE/S | IM mRNA; IN NE/IVT/S | IN NE/S; IN NE/S | IN NE/IVT/S; IN NE/IVT/S | IM PBS; IN NE/S | IM PBS; IN NE/IVT/S |
|---|---|---|---|---|---|---|---|---|---|
| IM mRNA; IM mRNA | | ns | ns | * | ** | ns | ** | ns | ns |
| IM mRNA; IN PBS | | | * | * | ** | ns | ** | ns | ns |
| IM mRNA; IN S only | | | | ns | ** | ns | * | ns | ns |
| IM mRNA; IN NE/S | | | | | ns | ns | ns | ns | ns |
| IM mRNA; IN NE/IVT/S | | | | | | ** | * | ** | * |
| IN NE/S; IN NE/S | | | | | | | ns | ns | ns |
| IN NE/IVT/S; IN NE/IVT/S | | | | | | | | ns | ns |
| IM PBS; IN NE/S | | | | | | | | | ns |
| IM PBS; IN NE/IVT/S | | | | | | | | | |

**Figure 4D**: cLN TNFa

| | IM mRNA; IM mRNA | IM mRNA; IN PBS | IM mRNA; IN S only | IM mRNA; IN NE/S | IM mRNA; IN NE/IVT/S | IN NE/S; IN NE/S | IN NE/IVT/S; IN NE/IVT/S | IM PBS; IN NE/S | IM PBS; IN NE/IVT/S |
|---|---|---|---|---|---|---|---|---|---|
| IM mRNA; IM mRNA | | ns | ns | ns | ns | ns | ns | ns | * |
| IM mRNA; IN PBS | | | ns | * | ** | ns | ** | ns | * |
| IM mRNA; IN S only | | | | ns | ** | ns | * | ns | * |
| IM mRNA; IN NE/S | | | | | ns | ns | ns | ns | * |
| IM mRNA; IN NE/IVT/S | | | | | | ** | ns | ** | ns |
| IN NE/S; IN NE/S | | | | | | | * | ns | * |
| IN NE/IVT/S; IN NE/IVT/S | | | | | | | | * | ns |
| IM PBS; IN NE/S | | | | | | | | | * |
| IM PBS; IN NE/IVT/S | | | | | | | | | |

**Figure 4E**: cLN IL5

| | IM mRNA; IM mRNA | IM mRNA; IN PBS | IM mRNA; IN S only | IM mRNA; IN NE/S | IM mRNA; IN NE/IVT/S | IN NE/S; IN NE/S | IN NE/IVT/S; IN NE/IVT/S | IM PBS; IN NE/S | IM PBS; IN NE/IVT/S |
|---|---|---|---|---|---|---|---|---|---|
| IM mRNA; IM mRNA | | ns | ns | ns | ns | ns | ns | ns | ns |
| IM mRNA; IN PBS | | | ns | ns | ns | ns | ns | ns | ns |
| IM mRNA; IN S only | | | | ns | ns | ns | ns | ns | ns |
| IM mRNA; IN NE/S | | | | | ns | ns | ns | ns | ns |
| IM mRNA; IN NE/IVT/S | | | | | | ns | ns | ** | * |
| IN NE/S; IN NE/S | | | | | | | ns | ns | ns |
| IN NE/IVT/S; IN NE/IVT/S | | | | | | | | * | ns |
| IM PBS; IN NE/S | | | | | | | | | ns |
| IM PBS; IN NE/IVT/S | | | | | | | | | |

**Figure 4F**: cLN IL13

| | IM mRNA; IM mRNA | IM mRNA; IN PBS | IM mRNA; IN S only | IM mRNA; IN NE/S | IM mRNA; IN NE/IVT/S | IN NE/S; IN NE/S | IN NE/IVT/S; IN NE/IVT/S | IM PBS; IN NE/S | IM PBS; IN NE/IVT/S |
|---|---|---|---|---|---|---|---|---|---|
| IM mRNA; IM mRNA | | ns | ns | ns | ns | ns | ns | ns | ns |
| IM mRNA; IN PBS | | | ns | ns | * | ns | * | ns | ns |
| IM mRNA; IN S only | | | | ns | ns | ns | ns | * | ns |
| IM mRNA; IN NE/S | | | | | ns | ns | ns | ns | ns |
| IM mRNA; IN NE/IVT/S | | | | | | ns | ns | ** | ns |
| IN NE/S; IN NE/S | | | | | | | ns | ns | ns |
| IN NE/IVT/S; IN NE/IVT/S | | | | | | | | * | ns |
| IM PBS; IN NE/S | | | | | | | | | ns |
| IM PBS; IN NE/IVT/S | | | | | | | | | |

**Figure 4G**: cLN IL6

| | IM mRNA; IM mRNA | IM mRNA; IN PBS | IM mRNA; IN S only | IM mRNA; IN NE/S | IM mRNA; IN NE/IVT/S | IN NE/S; IN NE/S | IN NE/IVT/S; IN NE/IVT/S | IM PBS; IN NE/S | IM PBS; IN NE/IVT/S |
|---|---|---|---|---|---|---|---|---|---|
| IM mRNA; IM mRNA | | ns | ns | ns | ns | ns | ns | ns | ns |
| IM mRNA; IN PBS | | | ns | ** | ** | ns | * | ns | * |
| IM mRNA; IN S only | | | | ** | ** | ns | * | ns | * |
| IM mRNA; IN NE/S | | | | | ns | ns | ns | ns | ns |
| IM mRNA; IN NE/IVT/S | | | | | | ** | ns | ns | ns |
| IN NE/S; IN NE/S | | | | | | | ns | ns | ns |
| IN NE/IVT/S; IN NE/IVT/S | | | | | | | | ns | ns |
| IM PBS; IN NE/S | | | | | | | | | ns |
| IM PBS; IN NE/IVT/S | | | | | | | | | |

**Figure 4H**: cLN IL17A

| | IM mRNA; IM mRNA | IM mRNA; IN PBS | IM mRNA; IN S only | IM mRNA; IN NE/S | IM mRNA; IN NE/IVT/S | IN NE/S; IN NE/S | IN NE/IVT/S; IN NE/IVT/S | IM PBS; IN NE/S | IM PBS; IN NE/IVT/S |
|---|---|---|---|---|---|---|---|---|---|
| IM mRNA; IM mRNA | | ns | * | ** | ** | ** | ** | ** | * |
| IM mRNA; IN PBS | | | ** | ** | ** | ** | ** | ** | * |
| IM mRNA; IN S only | | | | * | ** | ** | ** | ns | ns |
| IM mRNA; IN NE/S | | | | | ** | ns | ** | ns | ns |
| IM mRNA; IN NE/IVT/S | | | | | | ns | ns | ns | ns |
| IN NE/S; IN NE/S | | | | | | | ns | ns | ns |
| IN NE/IVT/S; IN NE/IVT/S | | | | | | | | * | ns |
| IM PBS; IN NE/S | | | | | | | | | ns |
| IM PBS; IN NE/IVT/S | | | | | | | | | |

**Figure 4I**: cLN IL10

| | IM mRNA; IM mRNA | IM mRNA; IN PBS | IM mRNA; IN S only | IM mRNA; IN NE/S | IM mRNA; IN NE/IVT/S | IN NE/S; IN NE/S | IN NE/IVT/S; IN NE/IVT/S | IM PBS; IN NE/S | IM PBS; IN NE/IVT/S |
|---|---|---|---|---|---|---|---|---|---|
| IM mRNA; IM mRNA | | ns | ns | ns | * | ns | * | ns | ns |
| IM mRNA; IN PBS | | | ns | ** | ** | ** | ** | ns | * |
| IM mRNA; IN S only | | | | ** | ** | ns | ** | ns | ns |
| IM mRNA; IN NE/S | | | | | ns | * | ns | * | * |
| IM mRNA; IN NE/IVT/S | | | | | | ** | ns | ** | * |
| IN NE/S; IN NE/S | | | | | | | ns | ns | ns |
| IN NE/IVT/S; IN NE/IVT/S | | | | | | | | * | * |
| IM PBS; IN NE/S | | | | | | | | | ns |
| IM PBS; IN NE/IVT/S | | | | | | | | | |

**Figure 6A:** 129S1 wk 6 serum IgG against full-length WT S

| | IM mRNA; IM mRNA | IM mRNA; IN S only | IM mRNA; IN NE/S | IM mRNA; IN NE/IVT/S | IN NE/S; IN NE/S | IN NE/IVT/S; IN NE/IVT/S | IM PBS; IN NE/S |
|---|---|---|---|---|---|---|---|
| IM mRNA; IM mRNA | | ns | ns | ns | ns | ns | ** |
| IM mRNA; IN S only | | | ** | ** | ns | ns | ** |
| IM mRNA; IN NE/S | | | | ns | ** | * | ** |
| IM mRNA; IN NE/IVT/S | | | | | ** | ** | ** |
| IN NE/S; IN NE/S | | | | | | ns | ** |
| IN NE/IVT/S; IN NE/IVT/S | | | | | | | ** |
| IM PBS; IN NE/S | | | | | | | |
| IM PBS; IN NE/IVT/S | | | | | | | |
| IM Advx/S; IM Advx/S | | | | | | | |
| IN PBS; IN PBS | | | | | | | |

**Figure 6B:** 129S1 wk6 serum WT MNT

| | IM mRNA; IM mRNA | IM mRNA; IN S only | IM mRNA; IN NE/S | IM mRNA; IN NE/IVT/S | IN NE/S; IN NE/S | IN NE/IVT/S; IN NE/IVT/S | IM PBS; IN NE/S |
|---|---|---|---|---|---|---|---|
| IM mRNA; IM mRNA | | ns | ns | ns | * | ns | ** |
| IM mRNA; IN S only | | | ns | ns | ns | ns | ** |
| IM mRNA; IN NE/S | | | | ns | * | ns | ** |
| IM mRNA; IN NE/IVT/S | | | | | * | ns | ** |
| IN NE/S; IN NE/S | | | | | | ns | ** |
| IN NE/IVT/S; IN NE/IVT/S | | | | | | | ** |
| IM PBS; IN NE/S | | | | | | | |
| IM PBS; IN NE/IVT/S | | | | | | | |
| IM Advx/S; IM Advx/S | | | | | | | |

**Figure 6C:** 129S1 wk6 serum B.1.351 MNT

| | IM mRNA; IM mRNA | IM mRNA; IN S only | IM mRNA; IN NE/S | IM mRNA; IN NE/IVT/S | IN NE/S; IN NE/S | IN NE/IVT/S; IN NE/IVT/S | IM PBS; IN NE/S |
|---|---|---|---|---|---|---|---|
| IM mRNA; IM mRNA | | ns | ns | ns | * | ns | * |
| IM mRNA; IN S only | | | ns | ns | ns | ns | ns |
| IM mRNA; IN NE/S | | | | ns | ns | ns | * |
| IM mRNA; IN NE/IVT/S | | | | | ns | ns | ns |
| IN NE/S; IN NE/S | | | | | | ns | ns |
| IN NE/IVT/S; IN NE/IVT/S | | | | | | | * |
| IM PBS; IN NE/S | | | | | | | |
| IM PBS; IN NE/IVT/S | | | | | | | |
| IM Advx/S; IM Advx/S | | | | | | | |

**Figure 6D:** 129S1 wk6 serum BA.1 MNT

| | IM mRNA; IM mRNA | IM mRNA; IN S only | IM mRNA; IN NE/S | IM mRNA; IN NE/IVT/S | IN NE/S; IN NE/S | IN NE/IVT/S; IN NE/IVT/S | IM PBS; IN NE/S |
|---|---|---|---|---|---|---|---|
| IM mRNA; IM mRNA | | ns | ns | ** | ns | ns | ns |
| IM mRNA; IN S only | | | ns | ** | ns | ns | ns |
| IM mRNA; IN NE/S | | | ns | ** | ns | ns | ns |
| IM mRNA; IN NE/IVT/S | | | | ns | ns | ns | * |
| IN NE/S; IN NE/S | | | | | | * | ** |
| IN NE/IVT/S; IN NE/IVT/S | | | | | | ns | ns |
| IM PBS; IN NE/S | | | | | | | ns |
| IM PBS; IN NE/IVT/S | | | | | | | |
| IM Advx/S; IM Advx/S | | | | | | | |

**Figure 6E:** 129S1 wk6 serum BA.4/5 MNT

| | IM mRNA; IM mRNA | IM mRNA; IN S only | IM mRNA; IN NE/S | IM mRNA; IN NE/IVT/S | IN NE/S; IN NE/S | IN NE/IVT/S; IN NE/IVT/S | IM PBS; IN NE/S |
|---|---|---|---|---|---|---|---|
| IM mRNA; IM mRNA | | ns | ns | ns | ns | ns | ns |
| IM mRNA; IN S only | | | ns | ns | ns | ns | * |
| IM mRNA; IN NE/S | | | | ns | ns | ns | * |
| IM mRNA; IN NE/IVT/S | | | | | ns | ns | * |
| IN NE/S; IN NE/S | | | | | | ns | * |
| IN NE/IVT/S; IN NE/IVT/S | | | | | | | * |
| IM PBS; IN NE/S | | | | | | | |
| IM PBS; IN NE/IVT/S | | | | | | | |
| IM Advx/S; IM Advx/S | | | | | | | |

**Figure 6F:** K18-hACE2 wk 6 serum IgG against full-length WT S

| | IM mRNA; IM mRNA | IM mRNA; IN PBS | IM mRNA; IN S only | IM mRNA; IN NE/S | IM mRNA; IN NE/IVT/S | IM Advx/S; IM Advx/S | IN PBS; IN PBS |
|---|---|---|---|---|---|---|---|
| IM mRNA; IM mRNA | | ** | ns | ns | ns | ns | ** |
| IM mRNA; IN PBS | | | ** | ** | * | ** | ** |
| IM mRNA; IN S only | | | | ns | ns | ns | ** |
| IM mRNA; IN NE/S | | | | | ns | ns | ** |
| IM mRNA; IN NE/IVT/S | | | | | | ns | ** |
| IM Advx/S; IM Advx/S | | | | | | | ** |
| IN PBS; IN PBS | | | | | | | |

**Figure 6G:** K18-hACE2 wk6 serum WT MNT

| | IM mRNA; IM mRNA | IM mRNA; IN PBS | IM mRNA; IN S only | IM mRNA; IN NE/S | IM mRNA; IN NE/IVT/S | IM Advx/S; IM Advx/S | IN PBS; IN PBS |
|---|---|---|---|---|---|---|---|
| IM mRNA; IM mRNA | | * | ns | ns | ns | ns | ** |
| IM mRNA; IN PBS | | | ns | * | ns | ns | ** |
| IM mRNA; IN S only | | | | ns | ns | ns | ** |
| IM mRNA; IN NE/S | | | | | ns | ns | ** |
| IM mRNA; IN NE/IVT/S | | | | | | ns | ** |
| IM Advx/S; IM Advx/S | | | | | | | ** |
| IN PBS; IN PBS | | | | | | | |

**Figure 6H:** K18-hACE2 wk6 serum B.1.351 MNT

| | IM mRNA; IM mRNA | IM mRNA; IN PBS | IM mRNA; IN S only | IM mRNA; IN NE/S | IM mRNA; IN NE/IVT/S | IM Advx/S; IM Advx/S | IN PBS; IN PBS |
|---|---|---|---|---|---|---|---|
| IM mRNA; IM mRNA | | ** | ns | ns | ns | ns | ** |
| IM mRNA; IN PBS | | | ** | ** | * | ns | ns |
| IM mRNA; IN S only | | | | ns | ns | ns | ** |
| IM mRNA; IN NE/S | | | | | ns | ns | ** |
| IM mRNA; IN NE/IVT/S | | | | | | ns | ** |
| IM Advx/S; IM Advx/S | | | | | | | * |
| IN PBS; IN PBS | | | | | | | |

**Figure 6I:** K18-hACE2 wk6 serum BA.1 MNT

| | IM mRNA; IM mRNA | IM mRNA; IN PBS | IM mRNA; IN S only | IM mRNA; IN NE/S | IM mRNA; IN NE/IVT/S | IM Advx/S; IM Advx/S | IN PBS; IN PBS |
|---|---|---|---|---|---|---|---|
| IM mRNA; IM mRNA | | ** | ns | ns | ns | ns | ** |
| IM mRNA; IN PBS | | | ** | ** | * | ** | ns |
| IM mRNA; IN S only | | | | ns | ns | ns | ** |
| IM mRNA; IN NE/S | | | | | ns | ns | ** |
| IM mRNA; IN NE/IVT/S | | | | | | ns | ** |
| IM Advx/S; IM Advx/S | | | | | | | ** |
| IN PBS; IN PBS | | | | | | | |

**Figure 6J:** K18-hACE2 wk6 serum BA.4/5 MNT

| | IM mRNA; IM mRNA | IM mRNA; IN PBS | IM mRNA; IN S only | IM mRNA; IN NE/S | IM mRNA; IN NE/IVT/S | IM Advx/S; IM Advx/S | IN PBS; IN PBS |
|---|---|---|---|---|---|---|---|
| IM mRNA; IM mRNA | | ns | ns | ns | ns | ns | ns |
| IM mRNA; IN PBS | | | ns | ns | * | ns | ns |
| IM mRNA; IN S only | | | | ns | * | ns | ns |
| IM mRNA; IN NE/S | | | | | ns | ns | ns |
| IM mRNA; IN NE/IVT/S | | | | | | ns | ** |
| IM Advx/S; IM Advx/S | | | | | | | ns |
| IN PBS; IN PBS | | | | | | | |

| Figure 5: Spleen IFN-$\gamma^{+}$ | IM mRNA; IM mRNA | IM mRNA; IN PBS | IM mRNA; IN S | IM mRNA; IN NE/S | IM mRNA; IN NE/IVT/S | IN NE/S; IN NE/S | IN NE/IVT/S; IN NE/IVT/S | IM PBS; IN PBS |
|---|---|---|---|---|---|---|---|---|
| IM mRNA; IM mRNA | | **** | ** | **** | ** | **** | **** | **** |
| IM mRNA; IN PBS | | | * | ns | ** | ns | ns | ns |
| IM mRNA; IN S only | | | | ns | ns | ** | * | ** |
| IM mRNA; IN NE/S | | | | | ns | ns | ns | ns |
| IM mRNA; IN NE/IVT/S | | | | | | *** | ** | *** |
| IN NE/S; IN NE/S | | | | | | | ns | ns |
| IN NE/IVT/S; IN NE/IVT/S | | | | | | | | ns |
| IM PBS; IN PBS | | | | | | | | |

| Figure 5: Spleen IL-$2^{+}$ | IM mRNA; IM mRNA | IM mRNA; IN PBS | IM mRNA; IN S only | IM mRNA; IN NE/S | IM mRNA; IN NE/IVT/S | IN NE/S; IN NE/S | IN NE/IVT/S; IN NE/IVT/S | IM PBS; IN PBS |
|---|---|---|---|---|---|---|---|---|
| IM mRNA; IM mRNA | | **** | ** | **** | ns | **** | **** | **** |
| IM mRNA; IN PBS | | | *** | ** | **** | ns | ns | ns |
| IM mRNA; IN S only | | | | ns | ns | ** | * | **** |
| IM mRNA; IN NE/S | | | | | ns | * | ns | *** |
| IM mRNA; IN NE/IVT/S | | | | | | **** | *** | **** |
| IN NE/S; IN NE/S | | | | | | | ns | ns |
| IN NE/IVT/S; IN NE/IVT/S | | | | | | | | ns |
| IM PBS; IN PBS | | | | | | | | |

| Figure 5: Spleen TNFa$^{+}$ | IM mRNA; IM mRNA | IM mRNA; IN PBS | IM mRNA; IN S only | IM mRNA; IN NE/S | IM mRNA; IN NE/IVT/S | IN NE/S; IN NE/S | IN NE/IVT/S; IN NE/IVT/S | IM PBS; IN PBS |
|---|---|---|---|---|---|---|---|---|
| IM mRNA; IM mRNA | | **** | ** | ** | * | **** | **** | **** |
| IM mRNA; IN PBS | | | ** | ** | *** | ns | ns | ns |
| IM mRNA; IN S only | | | | ns | ns | ** | * | *** |
| IM mRNA; IN NE/S | | | | | ns | ** | ns | *** |
| IM mRNA; IN NE/IVT/S | | | | | | *** | ** | **** |
| IN NE/S; IN NE/S | | | | | | | ns | ns |
| IN NE/IVT/S; IN NE/IVT/S | | | | | | | | ns |
| IM PBS; IN PBS | | | | | | | | |

| Figure 5: Spleen IFN-$\gamma^{+}$, IL-$2^{+}$, TNFa$^{+}$ | IM mRNA; IM mRNA | IM mRNA; IN PBS | IM mRNA; IN S only | IM mRNA; IN NE/S | IM mRNA; IN NE/IVT/S | IN NE/S; IN NE/S | IN NE/IVT/S; IN NE/IVT/S | IM PBS; IN PBS |
|---|---|---|---|---|---|---|---|---|
| IM mRNA; IM mRNA | | **** | * | ** | ns | **** | **** | **** |
| IM mRNA; IN PBS | | | * | ns | ** | ns | ns | ns |
| IM mRNA; IN S only | | | | ns | ns | ** | ** | ** |
| IM mRNA; IN NE/S | | | | | ns | * | * | * |
| IM mRNA; IN NE/IVT/S | | | | | | *** | *** | *** |
| IN NE/S; IN NE/S | | | | | | | ns | ns |
| IN NE/IVT/S; IN NE/IVT/S | | | | | | | | ns |
| IM PBS; IN PBS | | | | | | | | |

| Figure 5: cLN IFN-$\gamma^{+}$ | IM mRNA; IM mRNA | IM mRNA; IN PBS | IM mRNA; IN S | IM mRNA; IN NE/S | IM mRNA; IN NE/IVT/S | IN NE/S; IN NE/S | IN NE/IVT/S; IN NE/IVT/S | IM PBS; IN PBS |
|---|---|---|---|---|---|---|---|---|
| IM mRNA; IM mRNA | | ns | ns | * | *** | ns | ns | ns |
| IM mRNA; IN PBS | | | ns | *** | **** | ns | ns | ns |
| IM mRNA; IN S only | | | | * | *** | ns | ns | ns |
| IM mRNA; IN NE/S | | | | | ns | ** | *** | *** |
| IM mRNA; IN NE/IVT/S | | | | | | **** | **** | **** |
| IN NE/S; IN NE/S | | | | | | | ns | ns |
| IN NE/IVT/S; IN NE/IVT/S | | | | | | | | ns |
| IM PBS; IN PBS | | | | | | | | |

| Figure 5: cLN IL-$2^{+}$ | IM mRNA; IM mRNA | IM mRNA; IN PBS | IM mRNA; IN S only | IM mRNA; IN NE/S | IM mRNA; IN NE/IVT/S | IN NE/S; IN NE/S | IN NE/IVT/S; IN NE/IVT/S | IM PBS; IN PBS |
|---|---|---|---|---|---|---|---|---|
| IM mRNA; IM mRNA | | ns | ns | **** | **** | ns | ** | ** |
| IM mRNA; IN PBS | | | ns | **** | **** | ** | **** | ns |
| IM mRNA; IN S only | | | | **** | **** | ns | *** | ns |
| IM mRNA; IN NE/S | | | | | ns | *** | ns | **** |
| IM mRNA; IN NE/IVT/S | | | | | | **** | * | **** |
| IN NE/S; IN NE/S | | | | | | | * | *** |
| IN NE/IVT/S; IN NE/IVT/S | | | | | | | | **** |
| IM PBS; IN PBS | | | | | | | | |

| Figure 5: cLN TNFa$^{+}$ | IM mRNA; IM mRNA | IM mRNA; IN PBS | IM mRNA; IN S only | IM mRNA; IN NE/S | IM mRNA; IN NE/IVT/S | IN NE/S; IN NE/S | IN NE/IVT/S; IN NE/IVT/S | IM PBS; IN PBS |
|---|---|---|---|---|---|---|---|---|
| IM mRNA; IM mRNA | | ns | ns | **** | **** | ns | * | ns |
| IM mRNA; IN PBS | | | ns | **** | **** | ns | *** | ns |
| IM mRNA; IN S only | | | | **** | **** | ns | * | ns |
| IM mRNA; IN NE/S | | | | | ns | **** | ns | **** |
| IM mRNA; IN NE/IVT/S | | | | | | **** | * | **** |
| IN NE/S; IN NE/S | | | | | | | * | ns |
| IN NE/IVT/S; IN NE/IVT/S | | | | | | | | **** |
| IM PBS; IN PBS | | | | | | | | |

| Figure 5: cLN IFN-$\gamma^{+}$, IL-$2^{+}$, TNFa$^{+}$ | IM mRNA; IM mRNA | IM mRNA; IN PBS | IM mRNA; IN S only | IM mRNA; IN NE/S | IM mRNA; IN NE/IVT/S | IN NE/S; IN NE/S | IN NE/IVT/S; IN NE/IVT/S | IM PBS; IN PBS |
|---|---|---|---|---|---|---|---|---|
| IM mRNA; IM mRNA | | ** | ns | ns | * | ** | ** | ** |
| IM mRNA; IN PBS | | | ns | **** | **** | ns | ns | ns |
| IM mRNA; IN S only | | | | ** | **** | ns | ns | ns |
| IM mRNA; IN NE/S | | | | | ns | **** | **** | **** |
| IM mRNA; IN NE/IVT/S | | | | | | **** | **** | **** |
| IN NE/S; IN NE/S | | | | | | | ns | ns |
| IN NE/IVT/S; IN NE/IVT/S | | | | | | | | ns |
| IM PBS; IN PBS | | | | | | | | |

| Figure 5: Lung IFN-$\gamma^{+}$ | IM mRNA; IM mRNA | IM mRNA; IN PBS | IM mRNA; IN S | IM mRNA; IN NE/S | IM mRNA; IN NE/IVT/S | IN NE/S; IN NE/S | IN NE/IVT/S; IN NE/IVT/S | IM PBS; IN PBS |
|---|---|---|---|---|---|---|---|---|
| IM mRNA; IM mRNA | | ns | * | ns | ns | ns | ns | ns |
| IM mRNA; IN PBS | | | *** | ** | ns | ns | ns | ns |
| IM mRNA; IN S only | | | | ns | ns | *** | *** | *** |
| IM mRNA; IN NE/S | | | | | ns | *** | *** | *** |
| IM mRNA; IN NE/IVT/S | | | | | | * | * | * |
| IN NE/S; IN NE/S | | | | | | | ns | ns |
| IN NE/IVT/S; IN NE/IVT/S | | | | | | | | ns |
| IM PBS; IN PBS | | | | | | | | |

| Figure 5: Lung IL-$2^{+}$ | IM mRNA; IM mRNA | IM mRNA; IN PBS | IM mRNA; IN S only | IM mRNA; IN NE/S | IM mRNA; IN NE/IVT/S | IN NE/S; IN NE/S | IN NE/IVT/S; IN NE/IVT/S | IM PBS; IN PBS |
|---|---|---|---|---|---|---|---|---|
| IM mRNA; IM mRNA | | ns | * | ns | ns | ns | ns | ns |
| IM mRNA; IN PBS | | | *** | *** | ns | ns | * | ns |
| IM mRNA; IN S only | | | | ns | ns | ** | ns | **** |
| IM mRNA; IN NE/S | | | | | ns | * | ns | *** |
| IM mRNA; IN NE/IVT/S | | | | | | ns | ns | * |
| IN NE/S; IN NE/S | | | | | | | ns | ns |
| IN NE/IVT/S; IN NE/IVT/S | | | | | | | | * |
| IM PBS; IN PBS | | | | | | | | |

| Figure 5: LungTNFa$^{+}$ | IM mRNA; IM mRNA | IM mRNA; IN PBS | IM mRNA; IN S only | IM mRNA; IN NE/S | IM mRNA; IN NE/IVT/S | IN NE/S; IN NE/S | IN NE/IVT/S; IN NE/IVT/S | IM PBS; IN PBS |
|---|---|---|---|---|---|---|---|---|
| IM mRNA; IM mRNA | | ns | ** | ns | ns | ns | ns | ns |
| IM mRNA; IN PBS | | | *** | ** | ns | ns | ns | ns |
| IM mRNA; IN S only | | | | ns | ns | ** | ns | **** |
| IM mRNA; IN NE/S | | | | | ns | ns | ns | ** |
| IM mRNA; IN NE/IVT/S | | | | | | ns | ns | * |
| IN NE/S; IN NE/S | | | | | | | ns | ns |
| IN NE/IVT/S; IN NE/IVT/S | | | | | | | | ns |
| IM PBS; IN PBS | | | | | | | | |

| Figure 5: Lung IFN-$\gamma^{+}$, IL-$2^{+}$, TNFa$^{+}$ | IM mRNA; IM mRNA | IM mRNA; IN PBS | IM mRNA; IN S only | IM mRNA; IN NE/S | IM mRNA; IN NE/IVT/S | IN NE/S; IN NE/S | IN NE/IVT/S; IN NE/IVT/S | IM PBS; IN PBS |
|---|---|---|---|---|---|---|---|---|
| IM mRNA; IM mRNA | | ns | * | * | ns | ns | ns | ns |
| IM mRNA; IN PBS | | | *** | *** | ns | ns | ns | ns |
| IM mRNA; IN S only | | | | ns | ns | *** | *** | *** |
| IM mRNA; IN NE/S | | | | | ns | *** | *** | *** |
| IM mRNA; IN NE/IVT/S | | | | | | ns | ns | ns |
| IN NE/S; IN NE/S | | | | | | | ns | ns |
| IN NE/IVT/S; IN NE/IVT/S | | | | | | | | ns |
| IM PBS; IN PBS | | | | | | | | |

**Table S2. Antibodies utilized in flow cytometry.**

| Antibody | Clone | Source | Catalog |
|---|---|---|---|
| Pacific Blue Anti-Mouse CD3 | 17A2 | Biolegend | 100214 |
| BV605 Anti-Mouse CD4 | RM4-5 | Biolegend | 100548 |
| APC-Cy7 Anti-Mouse CD8a | 53-6.7 | Biolegend | 100714 |
| AF647 Anti-Mouse IFN-γ | XMG1.2 | Biolegend | 505814 |
| PE Anti-Mouse IL-2 | JES6-5H4 | Biolegend | 503808 |
| PE-Cy7 Anti-Mouse TNFα | MP6-XT22 | Biolegend | 506324 |
| BV785 Anti-Mouse IL-17A | TC11-18H10.1 | Biolegend | 506928 |
